# Protective effects of polyphenol-rich extracts against neurotoxicity elicited by paraquat or rotenone in cellular models of Parkinson’s disease

**DOI:** 10.1101/2023.04.26.538474

**Authors:** Mitali A. Tambe, Aurelie de Rus Jacquet, Katherine E. Strathearn, Gad G. Yousef, Mary H. Grace, Mario G. Ferruzzi, Qingli Wu, James E. Simon, Mary Ann Lila, Jean-Christophe Rochet

## Abstract

Parkinson’s disease (PD) is a neurodegenerative disorder involving motor symptoms caused by a loss of dopaminergic neurons in the substantia nigra region of the brain. Epidemiological evidence suggests that anthocyanin (ANC) intake is associated with a low risk of PD. Previously, we reported that extracts enriched with ANC and proanthocyanidins (PAC) suppressed dopaminergic neuron death elicited by the PD-related toxin rotenone in a primary midbrain culture model. Here, we characterized botanical extracts enriched with a mixed profile of polyphenols, as well as a set of purified polyphenolic standards, in terms of their ability to mitigate dopaminergic cell death in midbrain cultures exposed to another PD-related toxicant, paraquat (PQ), and we examined underlying neuroprotective mechanisms. Extracts prepared from blueberries, black currants, grape seeds, grape skin, mulberries, and plums, as well as several ANC, were found to rescue dopaminergic neuron loss in PQ-treated cultures. Comparison of a subset of ANC-rich extracts for the ability to mitigate neurotoxicity elicited by PQ versus rotenone revealed that a hibiscus or plum extract was only neuroprotective in cultures exposed to rotenone or PQ, respectively. Several extracts or compounds with the ability to protect against PQ neurotoxicity increased the activity of the antioxidant transcription factor Nrf2 in cultured astrocytes, and PQ-induced dopaminergic cell death was attenuated in Nrf2-expressing midbrain cultures. In other studies, we found that extracts prepared from hibiscus, grape skin, or purple basil (but not plums) rescued defects in O_2_ consumption in neuronal cells treated with rotenone. Collectively, these findings suggest that extracts enriched with certain combinations of ANC, PAC, stilbenes, and other polyphenols could potentially slow neurodegeneration in the brains of individuals exposed to PQ or rotenone by activating cellular antioxidant mechanisms and/or alleviating mitochondrial dysfunction.

## Introduction

Parkinson’s disease (PD) is a debilitating neurodegenerative disorder characterized by a loss of dopaminergic neurons in the *substantia nigra* of the midbrain region^1^. Additionally, surviving neurons accumulate cytosolic inclusions named Lewy bodies enriched with fibrillar forms of the presynaptic protein α-synuclein (aSyn)^2, 3^. Mutations in the gene encoding aSyn have been linked to cases of familial PD^4, 5^. Mitochondrial dysfunction and oxidative stress are thought to be key factors in PD pathogenesis^6^. In support of this idea, the brains of PD patients show a decrease in the activity of complex I, an enzyme of the mitochondrial electron transport chain. The leakage of electrons that occurs as a result of this complex I defect is hypothesized to trigger a build-up of reactive oxygen species (ROS), and these in turn may play a role in the formation of toxic aSyn oligomers involved in neurodegeneration^4, 7–9^.

Epidemiological data suggest that the risk of PD increases with exposure to two environmental pollutants: rotenone, a flavonoid derivative used as a pesticide, and paraquat (PQ), a bipyridyl derivative used as an herbicide^10^. Rotenone is a mitochondrial complex I inhibitor that disrupts the electron transport chain, thereby increasing mitochondrial oxidative stress^11^. Rats exposed systemically to rotenone exhibit evidence of ROS accumulation, dopaminergic cell death, and the presence of aSyn-positive, Lewy-like inclusions in the *substantia nigra*^12, 13^. In contrast to rotenone, PQ is not a potent complex I inhibitor (IC_50_ = 8.1 mM compared to 14 nM in the case of rotenone)^14^. Instead, PQ triggers oxidative stress through a REDOX cycling mechanism^15^. Namely, cellular enzymes convert PQ from the dicationic form to the monocationic form, which in turn reacts with oxygen to regenerate the dicationic form, resulting in the formation of superoxide radicals.

Polyphenols are naturally occurring molecules in plants and fruits that have been studied extensively for their potential therapeutic benefit in a number of diseases, including mild cognitive impairment^16, 17^ and various neurodegenerative disorders^18–20^. Our group and others have shown that polyphenols can alleviate neurotoxicity in preclinical models of PD, including flavonoids (a subclass of polyphenols) such as green tea polyphenols^21–23^, anthocyanins (ANC)^24–27^, and isoflavones^28–30^, as well as the phenolic acid derivative curcumin^31^ and stilbenes such as resveratrol and oxyresveratrol^24, 32^. Polyphenols have been shown to protect against oxidative stress in a variety of disease models by scavenging free radicals and up-regulating endogenous cytoprotective responses^33, 34^. Cellular mechanisms involved in polyphenol-mediated neuroprotection include activation of cellular antioxidant pathways, attenuation of mitochondrial dysfunction, and amelioration of glial activation^18, 20, 24–26, 28, 29, 35^.

Epidemiological data suggest that a high intake of extracts enriched in various flavonoids, including berry ANC (the glycosylated form of anthocyanidins) and PAC, is associated with lower risk of PD^36, 37^. Consistent with this idea, we found that various ANC- and PAC-rich extracts protected cultured dopaminergic neurons against rotenone toxicity to a greater extent than extracts enriched in other polyphenols, including phenolic acids (PA)^24^. In the study described herein, we characterized the same panel of extracts, as well as a new extract prepared from wild blueberries (BB) (**Table 1**), in terms of their ability to mitigate PQ neurotoxicity, and we examined the underlying neuroprotective mechanisms. Our results show that non-identical sets of extracts with overlapping polyphenol profiles have protective activity against PQ- versus rotenone-mediated neurotoxicity, suggesting that polyphenols (in particular, ANC, PAC, and stilbenes) mitigate dopaminergic cell death elicited by these two PD-related insults by activating different combinations of neuroprotective responses. Identifying mechanisms by which polyphenolic extracts reduce neurotoxicity elicited by rotenone or PQ is an important step towards developing tailored neuroprotective strategies to slow nigral degeneration in the brains of individuals exposed to either toxic agent.

**Table 1.** List of botanical extracts tested in this study.

| Extract | Botanical source | Major polyphenols<sup>a</sup> |
| --- | --- | --- |
| Black currant (BC) | *Ribes nigrum* | ANC, PAC, PA |
| Blueberry (BB) | *Vaccinium corymbosum* | ANC, PAC, PA |
| Wild BB | *Vaccinium angustifolium* | ANC, PAC, PA |
| Grape seed (GS) | *Vitis vinifera* | PAC, PA<sup>b</sup> |
| Grape skin (GSk) | *Vitis vinifera* | ANC, PAC, PA, S |
| Hibiscus | *Hibiscus sabdariffa* | ANC |
| Mulberry | *Morus alba* | S |
| Plum | *Prunus domestica* | ANC, PAC, PA |
| Purple basil (PB) | *Ocimum basilicum* | ANC, PA |

<sup>a</sup>ANC, anthocyanin; PAC, proanthocyanidin; PA, phenolic acid; S, stilbene
<sup>b</sup>Predominantly gallic acid

## Materials and Methods

### Materials

Unless otherwise stated, chemicals were obtained from Sigma Chemical Co. (St. Louis, MO). Dulbecco’s modified Eagle’s medium (DMEM), fetal bovine serum (FBS), penicillin/streptomycin, Lipofectamine 2000, Optimem, trypsin-EDTA, 4’, 6-diamidino-2- phenylindole (DAPI), and Tris-glycine Ready gels were obtained from Invitrogen (Carlsbad, CA). The dopaminergic N27 cell line was provided by Dr. Curt Freed (University of Colorado). Human iPSC-derived astrocytes (iCell astrocytes) were obtained from Cellular Dynamics International (CDI) (Madison, WI). Pure malvidin-3-glucoside (M3G) and cyanidin-3-sophoroside (C3So) were purchased from Carbomer Inc. (San Diego, CA).

### Antibodies

The following antibodies were used in this study: chicken anti-MAP2 (catalog number CPCA- MAP2, EnCor Biotechnology, Gainesville, FL); rabbit anti-tyrosine hydroxylase (TH) (catalog number AB152, Millipore, Billerica, MA); and anti-rabbit IgG-Alexa Fluor 488 and anti-chicken IgG-Alexa Fluor 594 (Invitrogen, Carlsbad, CA).

### Preparation of botanical extracts

Anthocyanin and proanthocyanidin-enriched extracts were prepared from cultivated BB (*Vaccinium corymbosum*) (highbush), wild BB (*Vaccinium angustifolium*) (lowbush), plums (*Prunus domestica*), blackcurrants (BC) (*Ribes nigrum*), grape skin (*Vitis vinifera*), and mulberry (*Morus alba*) as described^24^. The BB extracts in particular were analyzed by HPLC and LC-MS to accurately quantify ANC and other diverse flavonoids characteristic of BB. PAC were separated according to their degree of polymerization by normal phase-HPLC with fluorescence detection, and peaks were quantified based on peak area measurements with reference to a standard curve constructed with a procyanidin B2 dimer. Total phenolic content was measured using the Folin-Ciocalteu assay, and results were expressed as gallic acid equivalent (GAE) as described^24^.

A grape seed (GS) extract (*Vitis vinifera*) was obtained from iBioCeuticals LLC (Eastham, MA)^24^. An extract was prepared from hibiscus (*Hibiscus sabdariffa*) using a ‘dark red’ source of the variety Vimto as described^24, 38^. A purple basil (PB) extract (*Ocimum basilicum*) was prepared from field-grown basil variety Red Rubin (Johnny’s Selected Seeds, Albion, ME) at the Rutgers Agricultural Experiment Station (Pittstown, NJ)^38–40^.

For cell culture experiments, the hibiscus extract, resveratrol, M3G, and C3So were dissolved in dimethyl sulfoxide (DMSO). The purple basil and mulberry extracts were dissolved in 50% DMSO/ethanol (v/v). The plum extract was dissolved in 30% (v/v) ethanol. All other extracts were dissolved in water.

### Preparation of rat primary cultures

Primary cultures were prepared via dissection of E17 embryos obtained from pregnant Sprague– Dawley rats (Harlan, Indianapolis, IN) as described^24–26, 29^. All animal handling procedures were approved by the Purdue Animal Care and Use Committee. To prepare mesencephalic cultures, the midbrain region containing the *substantia nigra* and ventral tegmental area was isolated stereoscopically, and the cells were dissociated with trypsin (final concentration, 26 μg/mL in 0.9% [w/v] NaCl). For experiments that involved analysis of neuroprotective activity, the dissociated cells were plated in a 48-well plate (pretreated with poly-L-lysine, 5 μg/mL) at a density of 163,500 cells per well in midbrain culture media, consisting of DMEM, 10% (v/v) FBS, 10% (v/v) horse serum, penicillin (10 U/mL), and streptomycin (10 μg/mL). Five days after plating, the cells were treated with cytosine arabinofuranoside (AraC) (20 µM, 48 h) to inhibit the proliferation of glial cells. At this stage (7 days in vitro (DIV)), ∼10-20% of the total cell population consisted of neurons that appeared differentiated with extended processes. For experiments that involved monitoring the activation of the Nrf2 pathway in midbrain astrocytes, the dissociated cells were plated in a 96-well, black clear-bottom plate (pretreated with poly-L-lysine, 10 μg/mL) at a density of 81,750 cells per well in midbrain culture media. After 5 DIV, the cultures were treated with AraC (20 μM, 72 h) before initiating experimental treatments.

To prepare enriched astrocytic cultures, the cortical region of brains from E17 rat embryos was isolated stereoscopically, and the cells were dissociated with trypsin (final concentration, 26 μg/mL in 0.9% [w/v] NaCl). The cells were plated at a density of ∼5 million cells in a 100 cm^2^ flask (pretreated with rat collagen, 25 μg/mL) in media consisting of DMEM, 10% (v/v) FBS, 10% (v/v) horse serum, penicillin (10 U/mL), and streptomycin (10 μg/mL). Two days after plating, the flask contained attached clusters of cells (mostly astrocytes with few neurons) and unattached cells (mostly neurons). To obtain a highly enriched population of astrocytes, the conditioned media was removed and replaced with fresh media. The media was replaced every two days until most of the astrocytes had spread out on the plate (generally after 7 DIV). The astrocyte-rich culture was passaged once and then plated in a 96-well, black clear-bottom plate (pretreated with poly-L-lysine, 10 μg/mL) at a density of 5,000 cells per well before being used for experiments.

### Analysis of neuroprotective activities of extracts and compounds

Neuroprotective activities of botanical extracts and compounds were assessed as described previously^24–26, 29^. Briefly, primary midbrain cultures (7 DIV) were incubated in the presence of extract or compound (or the corresponding vehicle) for 72 h. Next, the cultures were incubated in fresh media containing PQ (2.5 μM) or rotenone (25 nM) plus extract, compound, or vehicle for an additional 24 h. Control cultures were incubated in media without PQ, extract, or compound. The cultures were fixed, permeabilized, blocked, and incubated with primary antibodies specific for MAP2 (chicken, 1:2000) and TH (rabbit, 1:500) for 48 h at 4°C. After washing with PBS (10 mM Na_2_HPO_4_, 1.76 mM KH_2_PO_4_, 136 mM NaCl, 2.7 mM KCl, pH 7.4), the cells were incubated with a goat anti-chicken antibody conjugated to Alexa Fluor 594 and a goat anti-rabbit antibody conjugated to Alexa Fluor 488 (each at 1:1000) for 1 h at 22 °C. Prolong gold antifade reagent with DAPI was applied to each culture well, before sealing with a coverslip.

Relative dopaminergic cell viability was assessed by counting MAP2- and TH- immunoreactive neurons using a Nikon TE2000-U inverted fluorescence microscope (Nikon Instruments, Melville, NY) with a 20X objective. The investigator responsible for counting the cells was blinded to the sample identities. A minimum of 10 random fields of view were selected, and approximately 500 to 1000 MAP2^+^ neurons were counted per experiment for each treatment. Each experiment was conducted at least 3 times using embryonic cultures prepared from different pregnant rats. The data were expressed as the percentage of MAP2^+^ neurons that were also TH^+^ (this ratiometric approach was used to account for variations in cell plating density).

### Nrf2 transcriptional activity assay

An adenovirus encoding EGFP downstream of the SX2 (E1) enhancer and minimal promoter derived from the mouse heme oxygenase-1 (HO-1) gene^41–43^ was prepared and used to monitor activation of the Nrf2 pathway in cortical astrocytic cultures and primary midbrain cultures as described^25, 26, 29^. Cells plated in a 96-well, black clear-bottom plate (see above) were transduced with ARE-EGFP reporter virus at an MOI of 6.25 twenty-four hours after plating (astrocytes), or at an MOI of 10 seven days after plating (midbrain cultures). After 48 h, the cells were washed once with HBSS and incubated in fresh media supplemented with extract or vehicle for 24 h. Control cells were transduced with the ARE-EGFP virus for 48 h and then incubated in fresh media for another 24 h, in the absence of extract (negative control) or in the presence of curcumin (5 μM) (positive control). The cells were imaged for GFP fluorescence using an automated Cytation 3 Cell Imaging Reader (BioTek Instruments, Winooski, VT) equipped with a 4x objective. To quantify EGFP fluorescence, regions of interest (ROIs) were generated by the Gen5 2.05 data analysis software (BioTek) based on the cellular size range (20 to 400 µm) and a designated fluorescence intensity threshold. For each experiment, the threshold was adjusted so that the number of ROIs above the threshold in the curcumin-treated culture was 5- to 8-fold greater (or the overall EGFP fluorescence intensity was 1.5- to 2.5-fold greater) than that in the negative-control culture. Next, the cultures were fixed using 4% (v/v) PFA and incubated with the nuclear stain DAPI (300 nM) for 10 min at 37 °C. The cells were imaged for DAPI fluorescence using the Cytation 3 Cell Imaging Reader, and ROIs were generated by the Gen5 2.05 software based on a size range of 5 to 20 μm. The fluorescence intensity threshold was set so that most of the nuclei stained with DAPI were included among the detected ROIs. The number of ROIs for EGFP was divided by the total cell number (ROIs obtained from DAPI fluorescence) for each treatment and normalized to the control value to obtain a fold-change value.

Additional experiments were carried out with human iCell astrocytes produced at CDI (Madison, WI) by differentiating an iPSC line that was reprogrammed from fibroblasts obtained from an apparently healthy female individual without known PD-related mutations^25, 26, 29^. iCell astrocytes were plated at a density of 10,000 cells per well on a 96-well, black clear-bottom plate (pretreated with laminin, 10 μg/mL) in DMEM/F12, HEPES media supplemented with 2% (v/v) FBS and 1% (v/v) N-2 supplement. After 24 h, the cells were transduced with the ARE-EGFP reporter virus at an MOI of 25. The cells were incubated and analyzed as outlined above, except that after the 24 h incubation in fresh media, the cells were incubated in the presence of Hoechst nuclear stain (2 μg/mL in HBSS) for 15 min at 37 °C, washed in HBSS, and imaged for EGFP and Hoechst fluorescence in HBSS at 37 °C. To quantify Hoechst fluorescence, ROIs were generated by the software based on a size range of 10 to 40 μm. The fluorescence intensity threshold was set so that most of the nuclei stained with Hoechst were included among the detected ROIs. The number of ROIs for EGFP was divided by the total number of ROIs obtained from Hoechst fluorescence for each treatment and normalized to the control value.

### O_2_ consumption assay

N27 cells were grown in glucose-free RPMI 1640 media supplemented with 10% (v/v) FBS, 10 mM HEPES, 10 mM galactose, 2 mM glutamine, 1 mM sodium pyruvate, 100 U/mL penicillin, and 100 μg/mL streptomycin. The cells were plated on a 10 cm dish at a density of 500,000 cells/plate. After 24 h, the cells were treated with botanical extract or vehicle for 21 h and then incubated with 50 nM rotenone (with extract or vehicle) for 3 h. The cells were harvested via centrifugation at 700 x g (10 min, 4 °C) and resuspended in O_2_ consumption buffer (20 mM HEPES, pH 7.2). Cellular respiration was measured using a Clark-type oxygen electrode attached to a voltmeter (Digital Model 10 Controller, Rank Brothers, Ltd, Cambridge, UK). The electrode was allowed to stabilize in O_2_ consumption buffer at 37 °C for 30 min to ensure air saturation. To normalize the background current, the voltmeter was set to zero using a polarizing voltage of 0.60 V. An aliquot of 1.5 x 10^6^ cells was loaded into the respiration chamber, where the sensitivity control was set to 1 V. This setting corresponded to 100% of the O_2_ concentration (0.21 mM) in the air-saturated reaction medium before the start of respiration. The sample was continuously stirred at 840 rpm using a magnetic stir bar located inside the chamber. Using the Pico Technology software program (PicoTechnology, Ltd., Cambridgeshire, UK), the O_2_ level remaining in the chamber at any time during respiration was automatically logged (with 10 sec intervals) as a voltage, V_O2_, corresponding to the voltage generated by the reaction of O_2_ with the electrode. The voltage decreased progressively as O_2_ was consumed. Data were plotted as percent of O_2_ consumed after 500 sec.

### Statistical analysis

Data from measurements of primary neuron viability, ARE-EGFP fluorescence, and mitochondrial O_2_ consumption were analyzed via one-way ANOVA with Tukey’s multiple comparisons *post hoc* test using GraphPad Prism version 8.0 (La Jolla, CA). Prior to performing these ANOVA analyses, the data were subjected to a square root transformation (neuron viability data) or log transformation (fluorescence and O_2_ consumption data) to conform to ANOVA assumptions. For measurements of ARE-EGFP fluorescence, fold-change values were log-transformed, and the transformed data were analyzed using GraphPad Prism 8.0 via a one-sample t-test to determine whether the mean of the log(fold-change) was different from the hypothetical value of 0 (corresponding to a ratio of 1). The ‘n’ values specified in the figure legends represent the number of biological replicates (i.e. independent experiments involving cultures prepared at different times).

## Results

### Neuroprotective activities of ANC-rich extracts

In an initial set of experiments, we tested ANC-rich extracts prepared from BB (highbush), wild BB (lowbush), BC (*R. nigrum*), or PB (*O. basilicum*) for the ability to alleviate PQ neurotoxicity. The polyphenolic compositions of the BB, BC, and PB extracts were reported in an earlier paper from our group^24^, whereas the wild BB extract (prepared as a flavonoid-rich extract) was characterized in this study. HPLC analyses revealed that the wild BB extract contained 33.6 mg/g polyphenols, and the ANC:PAC:PA mass ratio was approximately 60:25:15. The distribution of ANC in the extract was similar to that determined for the highbush BB extract, and the ANC profiles for both extracts (as well as the PB extract^39^) were substantially more complex than that of the BC extract (**Supplementary Tables 1 and 2**)^24^.

To monitor the effects of the extracts on PQ-induced dopaminergic cell death, primary midbrain cultures were pre-treated with each extract or vehicle and then exposed to PQ. The cultures were stained for tyrosine hydroxylase (TH, a marker of dopaminergic neurons) and microtubule associated protein 2 (MAP2, a general neuronal marker), and relative dopaminergic cell survival was assessed by determining the percentage of MAP2^+^ neurons that also stained positive for TH. We found that cultures treated with PQ plus BB, wild BB, or BC extract (but not PB extract) had a higher percentage of dopaminergic neurons than cultures treated with PQ alone (**Figures 1A-E**). Additionally, the BB extract showed a trend towards inducing an increase in dopaminergic cell viability even when it was removed from the cultures 6 h prior to the addition of PQ, suggesting that the extract did not merely mitigate neurotoxicity by blocking the entry of PQ into the cells (**Figure 1B**). Collectively, these data suggest that some (but not all) ANC-rich extracts can interfere with PQ-mediated neuronal cell death in primary midbrain cultures. Moreover, the bioactivity of a given extract is dependent not only on the ANC concentration, but also on the types of phenolics in the extract.

**Figure 1.**
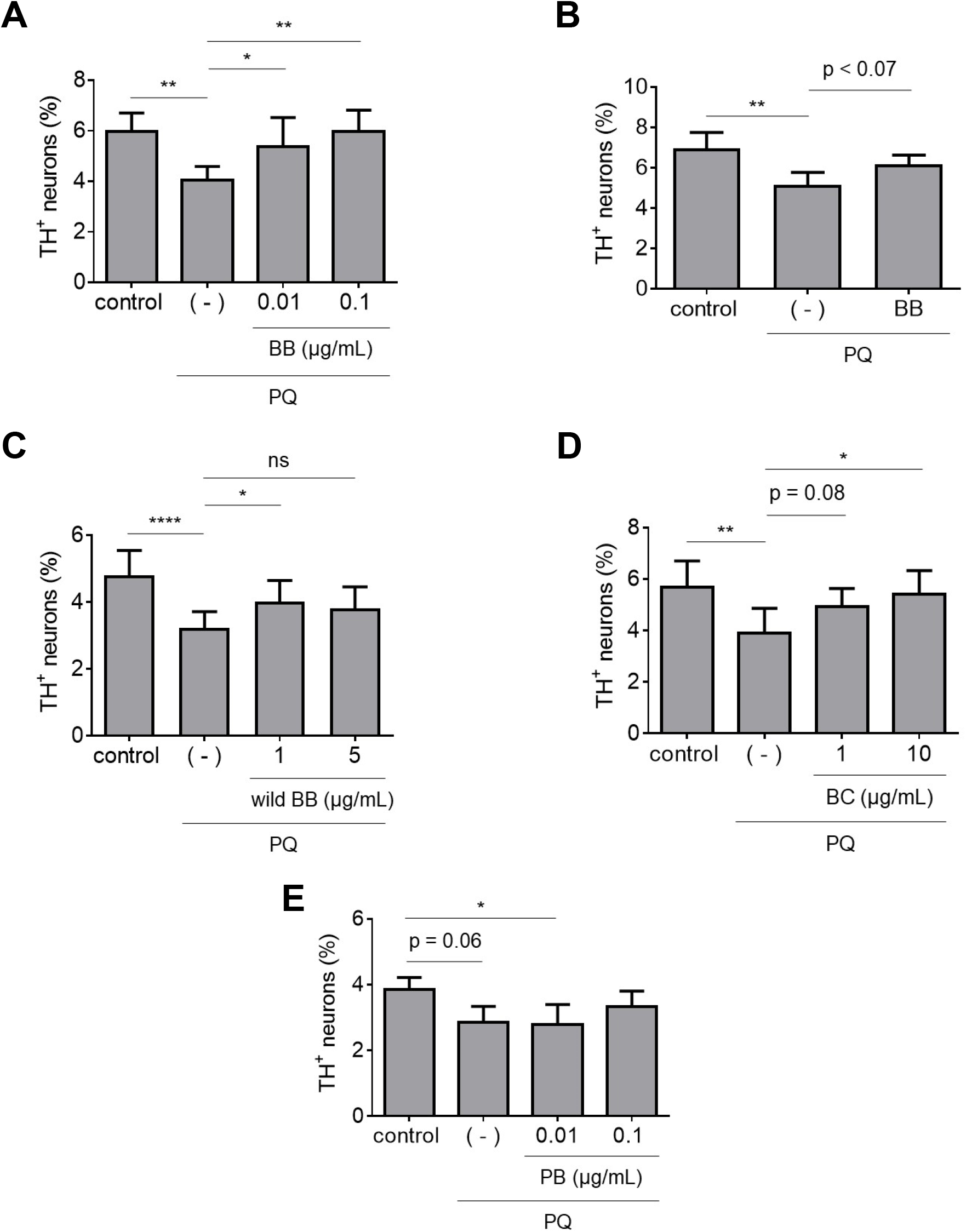
Polyphenol-enriched extracts with different ANC profiles have different abilities to protect against PQ neurotoxicity. Primary midbrain cultures incubated in the absence or presence of extract prepared from BB (A), wild BB (C), BC (D), or PB (E) for 72 h were exposed to PQ (2.5 μM) in the absence or presence of extract for 24 h. Alternatively, cultures incubated in the absence or presence of a BB extract (0.1 μg/mL) for 66 h were incubated in fresh media (minus extract) for 6 h and then exposed to PQ (2.5 μM) in the absence of extract for 24 h (B). Control cells were incubated in the absence of PQ or extract. The cells were stained with antibodies specific for MAP2 and TH and scored for relative dopaminergic cell viability. The data are presented as the mean ± SEM; *n* = 4 (A, D and E), *n* = 5 (B), or *n* = 6 (C); \**p*<0.05, \*\**p*<0.01, ****p<0.0001, square root transformation, one-way ANOVA with Tukey’s multiple comparisons post hoc test (ns, not significant). In panels (B), (D), and (E), a statistically significant neuroprotective effect is observed for cultures treated with BB (B) or BC (1 µg/mL; D) plus PQ versus PQ alone, and a significant neurotoxic effect is observed for cultures treated in the presence versus the absence of PQ (E), when the square root-transformed data are analyzed via ANOVA with the Newman–Keuls post hoc test (p ≤ 0.05).

### Neuroprotective activities of extracts rich in PAC and PA

Next, we tested a GS extract enriched in PAC and the PA gallic acid, as well as a plum extract with a polyphenol content of ∼65% PAC and ∼35% PA along with small amounts of ANC (**Supplementary Table 2**)^24^. We reasoned that comparing the neuroprotective activities of the GS and plum extracts should reveal the relative importance of PAC and PA in alleviating PQ neurotoxicity. An increase in dopaminergic neuron survival was observed in midbrain cultures treated with PQ plus either extract compared to cultures exposed to PQ alone (**Figures 2A,B**). To further evaluate the contribution of PA in the plum extract, we tested a PA derivative, curcumin. We found that the percentage of dopaminergic neurons in cultures exposed to PQ in the presence of curcumin was significantly higher than in cultures exposed to PQ alone (**Figure 2C**). Overall, these data suggest that PAC and PA have protective effects against PQ neurotoxicity.

**Figure 2.**
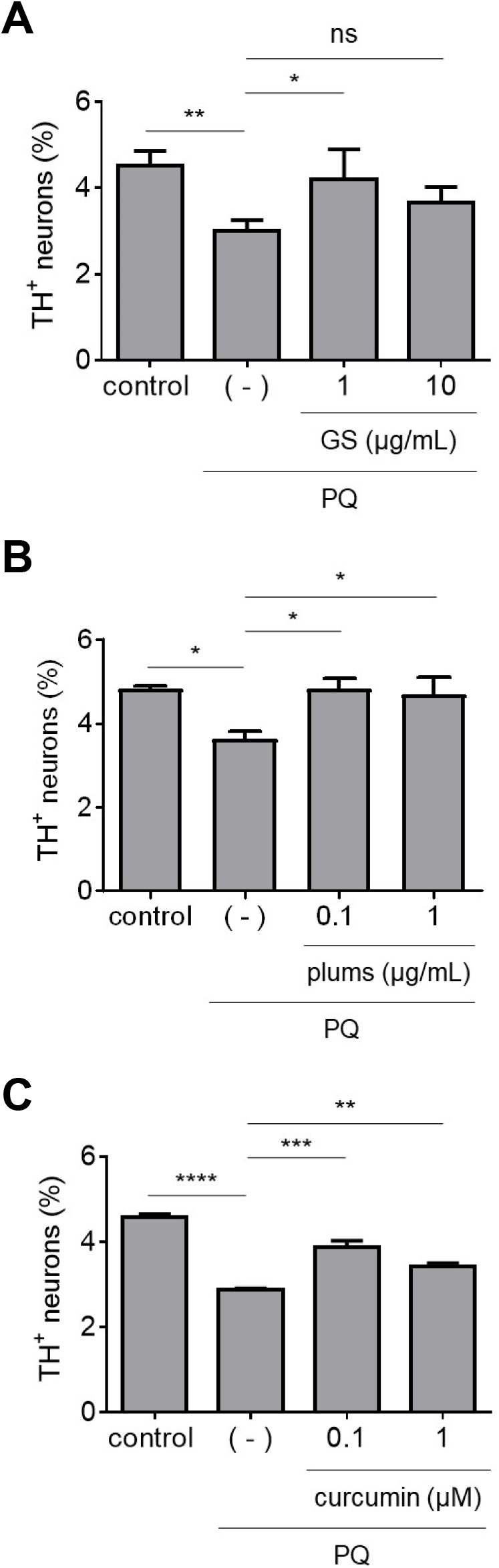
Extracts enriched with PAC or PA protect against PQ neurotoxicity. Primary midbrain cultures incubated in the absence or presence of a GS extract (A), plum extract (B), or curcumin (C) for 72 h were exposed to PQ (2.5 μM) in the absence or presence of extract or compound for 24 h. Control cells were incubated in the absence of PQ, extract, or compound. The cells were stained with antibodies specific for MAP2 and TH and scored for relative dopaminergic cell viability. The data are presented as the mean ± SEM; *n* = 3 (A and C) or *n* = 5 (B); \**p*<0.05, \*\**p*<0.01, \*\*\**p*<0.001, \*\*\*\**p*<0.0001, square root transformation, one-way ANOVA with Tukey’s multiple comparisons post hoc test (ns, not significant).

### Neuroprotective activities of stilbene-rich extracts

Stilbenes such as resveratrol and oxyresveratrol have been studied for their neuroprotective activities in preclinical models of neurodegenerative diseases^24, 32^. In this study, we tested a mulberry extract rich in oxyresveratrol and a grape skin (GSk) extract rich in resveratrol^24^, as well as pure resveratrol, for their ability to mitigate PQ-induced dopaminergic cell loss. We found that midbrain cultures treated with PQ plus the mulberry or GSk extract or pure resveratrol had a higher relative number of dopaminergic neurons compared to cultures treated with PQ plus vehicle (**Figures 3A-C**). Together, these data suggest that stilbene-rich extracts and resveratrol are protective against PQ neurotoxicity.

**Figure 3.**
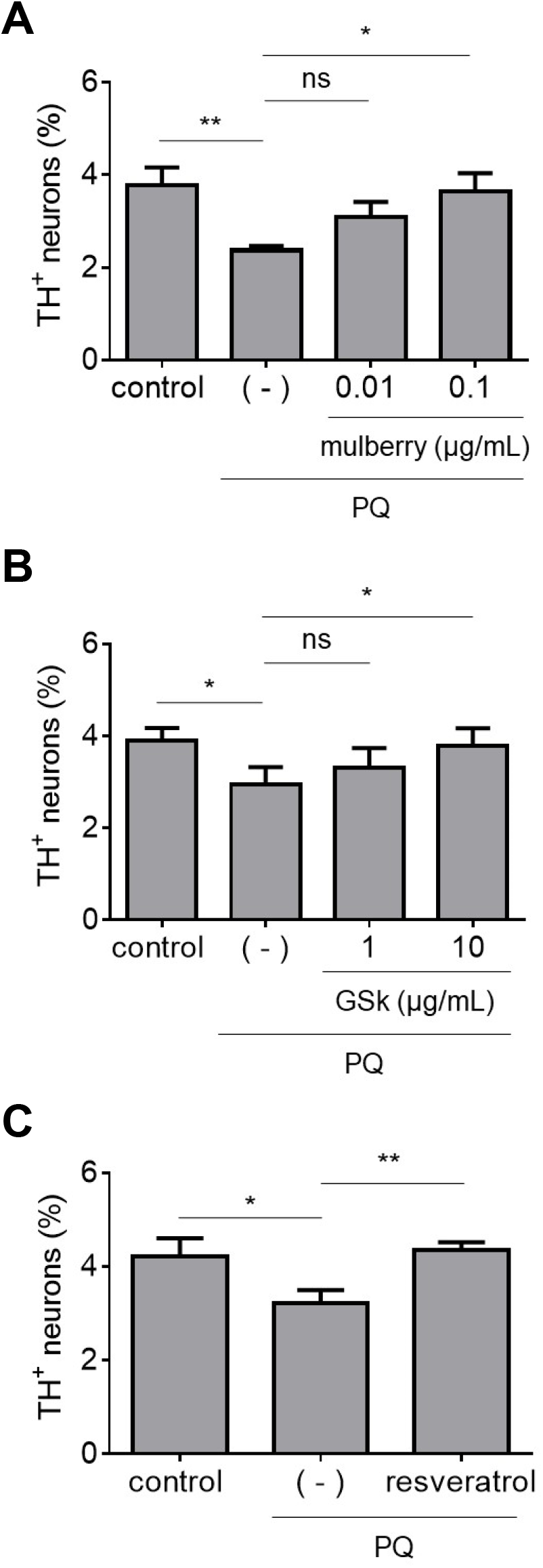
Stilbene-rich extracts alleviate PQ neurotoxicity. Primary midbrain cultures incubated in the absence or presence of a mulberry extract (A), GSk extract (B), or resveratrol (10 nM) (C) for 72 h were exposed to PQ (2.5 μM) in the absence or presence of extract or compound for 24 h. Control cells were incubated in the absence of PQ, extract, or compound. The cells were stained with antibodies specific for MAP2 and TH and scored for relative dopaminergic cell viability. The data are presented as the mean ± SEM; *n* = 4 (A and C) or *n* = 5 (B); \**p*<0.05, \*\**p*<0.01, square root transformation, one-way ANOVA with Tukey’s multiple comparisons post hoc test (ns, not significant).

### Neuroprotective activities of individual ANC

Because ANC-rich BB and BC extracts protected dopaminergic neurons against PQ neurotoxicity, we next tested the effects of individual ANC. We assessed two commercially available ANC, cyanidin-3-sophoroside (C3So) and malvidin-3-glucoside (M3G), and two ANC isolated from a BC extract by HPLC, cyanidin-3-glucoside (C3G) and delphinidin-3-glucoside (D3G), for the ability to protect against PQ neurotoxicity. We found that midbrain cultures treated with PQ plus C3G, D3G and M3G, but not C3So, exhibited higher dopaminergic cell viability than cultures treated with PQ alone (**Figures 4A-D**). Additionally, we tested a hibiscus extract that primarily consists of two ANC, cyanidin-3-O-sambubioside (C3Sa) and delphinidin-3-O-sambubioside (D3Sa)^24^. Interestingly, the hibiscus extract failed to attenuate dopaminergic cell death in primary midbrain cultures exposed to PQ (**Figure 4E**). Collectively, these data suggest that some (but not all) individual ANC can protect against PQ neurotoxicity.

**Figure 4.**
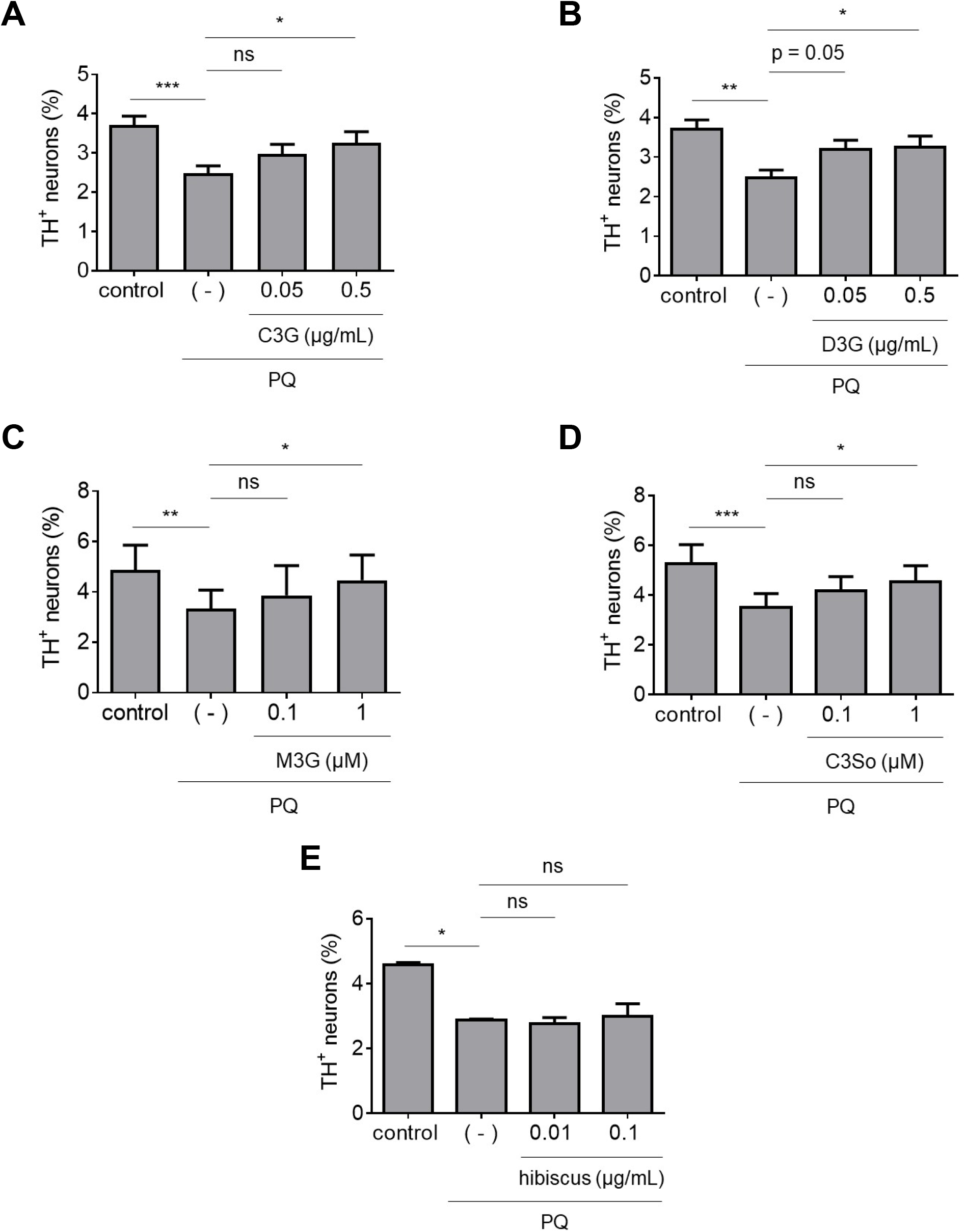
Individual ANC (but not a hibiscus extract) protect against PQ neurotoxicity. Primary midbrain cultures incubated in the absence or presence of C3G (A), D3G (B), M3G (C), C3So (D), or a hibiscus extract (E) for 72 h were exposed to PQ (2.5 μM) in the absence or presence of compound or extract for 24 h. Control cells were incubated in the absence of PQ, compound, or extract. The cells were stained with antibodies specific for MAP2 and TH and scored for relative dopaminergic cell viability. The data are presented as the mean ± SEM; *n* = 3 (E) or *n* = 4 (A, B and C) or *n* = 5 (D); \**p*<0.05, \*\**p*<0.01, ***p<0.001, square root transformation, one-way ANOVA with Tukey’s multiple comparisons post hoc test (ns, not significant).

### Selective protective activity of extracts against neurotoxicity elicited by PQ versus rotenone

In a previous study, we characterized the extracts examined here in terms of protective effects against neurotoxicity elicited by rotenone in primary midbrain cultures^24^. A comparison of the data obtained here and in our earlier study revealed that some extracts exhibited selective neuroprotective activity against toxicity elicited by PQ or rotenone. For example, the hibiscus extract alleviated dopaminergic cell death in cultures exposed to rotenone, but not PQ, whereas the plum extract protected against neurotoxicity elicited by PQ, but not rotenone. To confirm this selectivity in neuroprotective effects against different PD-related insults, we re-tested the hibiscus and plum extracts using the same cell culture preparations exposed to PQ or rotenone in parallel. The rotenone concentration used in this study was 25 nM as opposed to the concentration of 100 nM used in our previous study^24^ because we found that the lot of rotenone used here yielded a more consistent level of neurotoxicity at the lower concentration. The data showed that the plum extract mitigated neurotoxicity elicited by PQ, while only showing a trend towards protecting against dopaminergic neuron loss induced by rotenone, whereas the hibiscus extract mitigated dopaminergic cell death in cultures exposed to rotenone, but not PQ (**Supplementary Figure 1**). Collectively, these results indicate that some polyphenol-rich extracts have selective protective activity against neurotoxicity elicited by PQ versus rotenone.

### Effects of botanical extracts on Nrf2 transcriptional activity

Nrf2 is a redox-responsive transcription factor responsible for regulating the expression of cellular antioxidant proteins^44^. Multiple lines of evidence suggest that polyphenols modulate Nrf2 transcriptional activity^25, 26, 29, 45, 46^. Because PQ and rotenone are both thought to elicit neurotoxicity by eliciting oxidative stress, we hypothesized that at least a subset of botanical extracts might protect dopaminergic neurons against PD-related toxins by activating the Nrf2- mediated antioxidant response. Given that Nrf2 is predominantly expressed in astrocytes^47, 48^, we assessed the effects of botanical extracts on Nrf2 transcriptional activity in primary cortical astrocytes using an EGFP-based reporter assay. The assay involves transducing cells with adenovirus harboring a plasmid (pAd-ARE-EGFP-TKpolyA) encoding EGFP under the control of an enhancer with two AREs that serve as Nrf2 binding sites. Thus, an increase in Nrf2 transcriptional activity is detected as an increase in cellular EGFP fluorescence in the transduced cells. Transduced cortical astrocytes treated with curcumin (10 μM), a known Nrf2 activator that serves as a positive control in the reporter assay^25, 26, 29, 46^, showed a 5 to 8-fold increase in EGFP fluorescence intensity compared to control cells (data not shown). Cortical astrocytes treated with extract prepared from BB, wild BB, or GSk, or with the individual ANC C3G, D3G, or M3G, showed a ∼1.3 to 2.0-fold increase in EGFP fluorescence relative to control levels (**Figure 5**). Similar results were obtained in human iPSC-derived (iCell) astrocytes transduced with the reporter virus and treated with BB extract or M3G (**Supplementary Figure 2**). In contrast, an increase in EGFP fluorescence was not observed in cells treated with BC, PB, GS, plum, mulberry, or hibiscus extract (**Supplementary Figure 3**). Additionally, primary midbrain cultures transduced with Nrf2-encoding adenovirus at an MOI adjusted to achieve an ∼1.5-fold increase in Nrf2 transcriptional activity (as measured by the ARE-EGFP reporter assay; data not shown) showed a decrease in PQ-induced dopaminergic neuron death compared to untransduced cultures (**Figure 6**). Collectively, these data suggest that a subset of polyphenols in our panel of extracts alleviate PQ neurotoxicity by activating Nrf2-mediated transcription.

**Figure 5.**
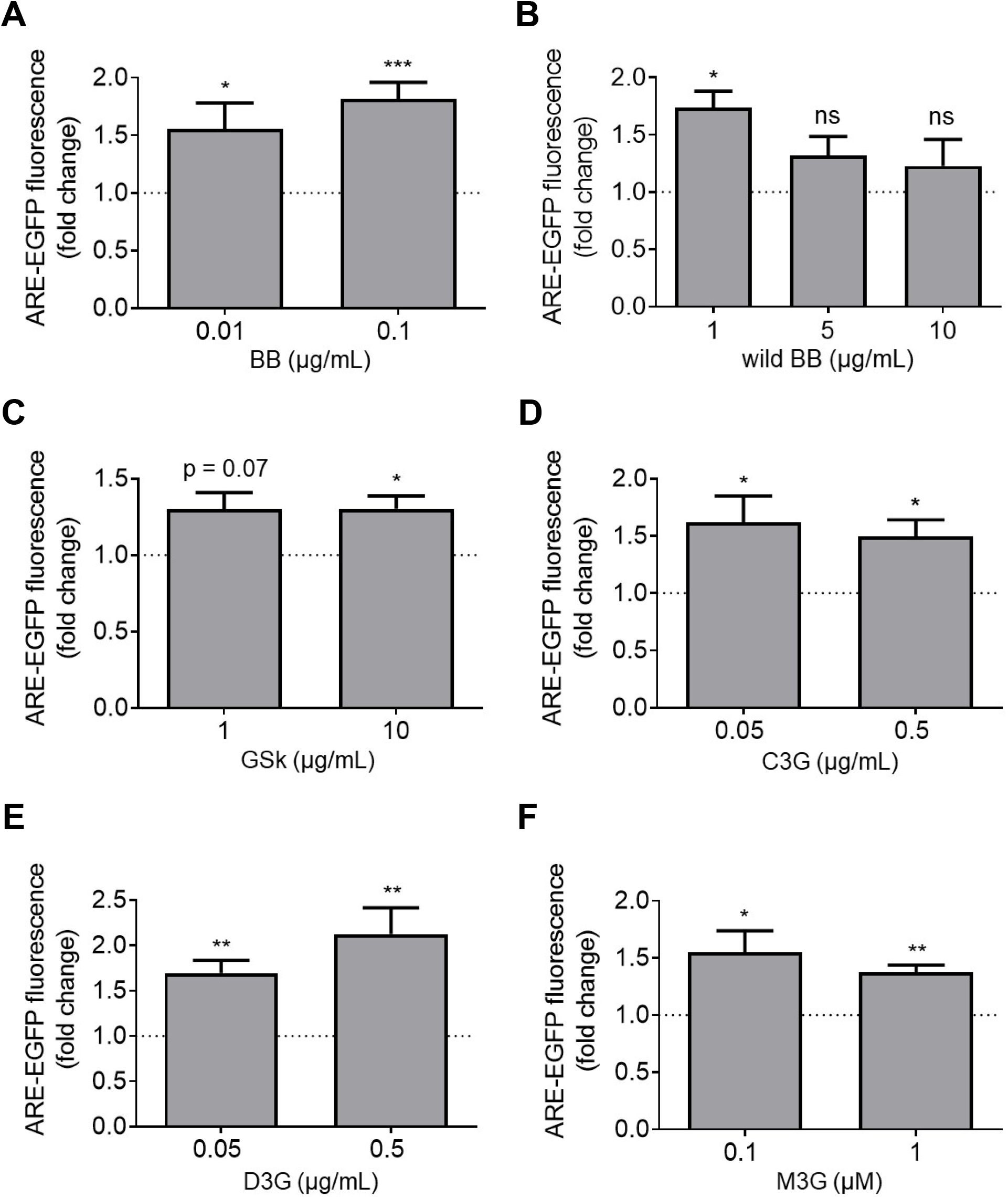
A subset of botanical extracts and all of the tested ANC increase Nrf2 transcriptional activity. Primary cortical astrocytes transduced with an ARE-EGFP reporter adenovirus for 48 h were incubated in the absence or presence of BB extract (A), wild BB extract (B), GSk extract (C), C3G (D), D3G (E), or M3G (F) for 24 h. Control astrocytes were transduced with the reporter virus and incubated in the absence of extract or compound. The cells were imaged to determine the intracellular EGFP fluorescence intensity. The data are presented as the mean ± SEM; *n* = 4 (B and C) or *n* = 7 (A, D, E and F); \**p*<0.05, \*\**p*<0.01, ***p<0.001 versus a predicted ratio of 1, log transformation followed by one-sample t-test (ns, not significant).

**Figure 6.**
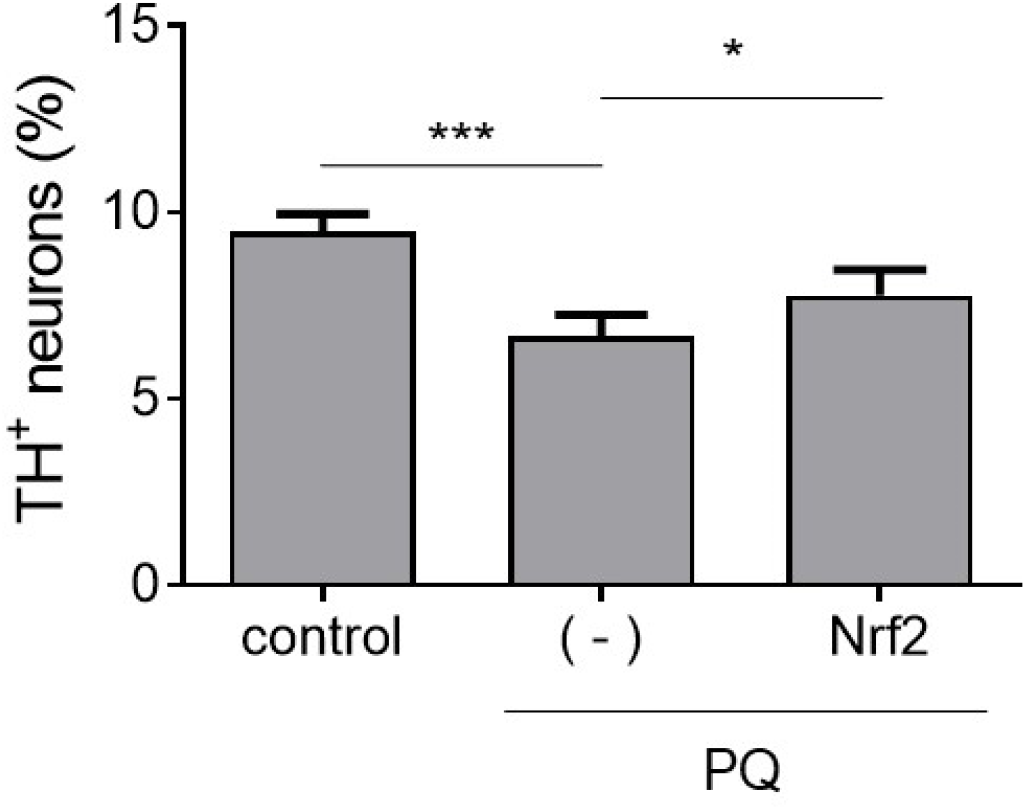
Overexpression of Nrf2 is sufficient for protection against PQ neurotoxicity. Primary midbrain cultures (untransduced or transduced with Nrf2 adenovirus) were incubated with PQ (2.5 µM). Control cells were incubated in the absence of virus or PQ. The cells were stained with antibodies specific for MAP2 and TH and scored for relative dopaminergic cell viability. The data are presented as the mean ± SEM; *n* = 3; \**p*<0.05, \*\*\**p*<0.001, square root transformation, one-way ANOVA with Tukey’s multiple comparisons post hoc test.

### Effects of botanical extracts on mitochondrial respiration

Next, we tested a subset of botanical extracts for the ability to rescue rotenone-induced defects in mitochondrial respiration. Data from several studies suggest that polyphenols and polyphenol-rich extracts can attenuate mitochondrial dysfunction^24, 26, 29, 49^. Rotenone elicits neurotoxicity by interfering with mitochondrial electron transport via complex I inhibition^11^, whereas PQ triggers a build-up of cytosolic ROS through a redox cycling mechanism^15^. Thus, PQ would be expected to have a less pronounced deleterious effect on mitochondrial function compared to rotenone, although a sufficiently high build-up of cytosolic ROS induced by PQ would be expected to ultimately cause mitochondrial dysfunction. Based on these considerations, we hypothesized that extracts with protective activity against rotenone neurotoxicity might rescue rotenone-mediated mitochondrial dysfunction. To address this hypothesis, we measured rates of cellular O_2_ consumption (a readout of mitochondrial respiration) using the N27 rat dopaminergic cell line. The cells were maintained in galactose media to avoid the ‘Crabtree effect’, a phenomenon wherein cells grown in high-glucose media bypass mitochondrial metabolism by using glycolysis as an alternative mode of energy production, thereby becoming resistant to toxicity elicited by inhibitors of mitochondrial function^26, 29, 50^. Galactose-conditioned N27 cells exhibited a pronounced loss of viability when exposed to rotenone at low nanomolar concentrations (similar to the concentration used in primary midbrain cultures), whereas cells cultured in high-glucose media were unaffected under these conditions (data not shown). Galactose-conditioned N27 cells treated with rotenone for 3 h showed a 50% reduction in cellular O_2_ consumption, and this decrease was mitigated by extracts prepared from PB, GSk, and hibiscus, but not plums (**Figure 7**). These data indicate that a subset of polyphenol-rich extracts that protect against rotenone neurotoxicity in primary midbrain cultures alleviate rotenone-induced mitochondrial dysfunction.

**Figure 7.**
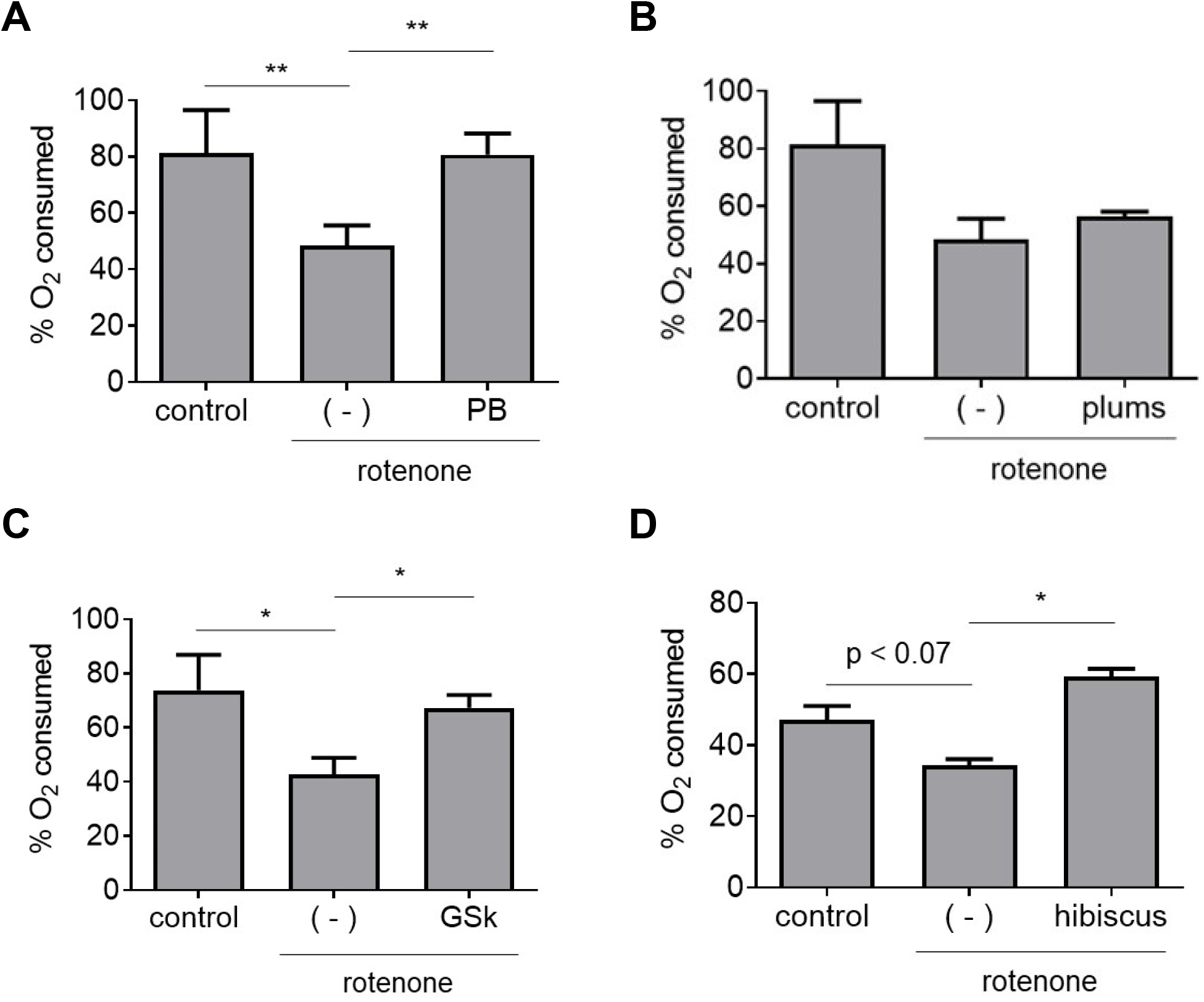
A subset of botanical extracts alleviate rotenone-induced deficits in mitochondrial respiration. Galactose-conditioned N27 cells were incubated in the absence or presence of an extract prepared from PB (A), plums (B), GSk (C) or hibiscus (D) for 21 h and then exposed to rotenone (50 nM) in the absence or presence of extract for 3 h. Control cell were incubated in the absence of rotenone or extract. Oxygen consumption was measured with a Clark-type oxygen electrode attached to a voltmeter. The data are presented as the mean ± SEM; *n* = 3 (A, C and D) or *n* = 4 (B); \**p*<0.05, \*\**p*<0.01, log transformation, one-way ANOVA with Tukey’s multiple comparisons post hoc test.

## Discussion

In this study, we characterized botanical extracts enriched in different polyphenols for their protective activity against PQ neurotoxicity in primary midbrain cultures. Because the cultures are composed of post-mitotic dopaminergic and non-dopaminergic neurons as well as glial cells, similar to the midbrain region affected in the brains of PD patients, they serve as an excellent cellular model of neurodegeneration in PD. Moreover, our approach of monitoring Nrf2 signaling in pure astrocytic cultures and O_2_ consumption in galactose-conditioned N27 dopaminergic neuronal cells enabled us to interrogate neuroprotective mechanisms in relevant cell models. By comparing the data obtained in this study with the results of a previous study involving a rotenone model of neurotoxicity^24^, we were able to identify polyphenol-rich extracts with neuroprotective activity against either or both PD-related insults.

### ANC- and PAC- rich extracts protect dopaminergic neurons against PQ neurotoxicity

An important outcome of this study was our finding that ANC-rich extracts prepared from BB, wild BB, and BC protected dopaminergic neurons against PQ toxicity. These results are consistent with previous data showing that an ANC-rich extract prepared from *Amelanchier arborea* by our group^25^ or from *Aronia melanocarpa* by Zimmerman and colleagues^51^ alleviates PQ neurotoxicity in cell culture. Here, we observed that the BB and wild BB extracts protected dopaminergic neurons against PQ toxicity at ∼10 to ∼1000-fold lower concentrations compared to the BC extract. This result is surprising given the markedly higher total ANC content of the BC extract^24^ but presumably can be attributed to the different ANC profiles of the three extracts. HPLC analysis revealed that the BB and wild BB extracts contained at least 17 and 18 different ANC, respectively, compared to only 4 ANC in the BC extract (**Supplementary Tables 1** and **2**). Synergism among different ANC^52^ could contribute to the more potent neuroprotective activity of the BB and wild BB extracts. Additionally, the two BB extracts (but not the BC extract) contained a considerable amount of PA (i.e., chlorogenic acid) that could also contribute to the extract’s neuroprotective activity.

In contrast to the BB, wild BB, and BC extracts, an ANC-rich PB extract failed to protect dopaminergic neurons against PQ neurotoxicity. Interestingly, the predominant ANC in the PB extract, malonyl and coumaryl glycoside derivatives of cyanidin and peonidin^39^, were absent from the BB, wild BB, and BC extracts (**Supplementary Tables 1 and 2**)^24^. Thus, we infer that differences in the protective activities of the BB, wild BB, BC, and PB extracts against PQ neurotoxicity could reflect differences in their total ANC content, ANC profiles, and/or distribution of other polyphenols (e.g. PAC and other flavonoids) that were below the detection limit of our RP-HPLC analysis^24^.

We also found that PAC-rich GS and plum extracts protected against PQ-induced neurotoxicity. Consistent with this observation, a PAC-rich extract prepared from the peels of the avocado *Persea americana* (var. Colinred) showed apparent neuroprotective effects (resulting in improved locomotor function and extended life span) in a *Drosophila* model of PQ exposure^53^. Data from other studies suggest that monomeric PACs including epicatechin and (-)- epigallocatechin-3-gallate can attenuate PQ-induced neurotoxicity in cell culture and *Drosophila* models^54, 55^. The relative *in vivo* potencies of the PAC-rich extracts examined here likely depend on their PAC profiles and the relative amounts/identities of monomeric versus polymeric PAC in the extracts, given evidence that (i) monomeric PAC have greater protective activity than polymeric PAC against toxicity elicited by PD-related insults^21, 56^, and (ii) monomeric PAC metabolites in their glucuronidated forms have greater brain bioavailability compared to polymeric _PAC_^57^, ^58^.

### Stilbene-rich extracts protect against PQ neurotoxicity

We observed that a mulberry extract rich in oxyresveratrol, a GSk extract rich in resveratrol, and pure resveratrol protected against PQ neurotoxicity. The concentration of resveratrol in the diluted GSk extract that showed neuroprotective activity (∼5 nM) was similar to that in the pure compound solution (10 nM)^24^, suggesting that resveratrol accounted for a major part of the extract’s ability to alleviate PQ-induced dopaminergic cell death. These results are consistent with previous data showing that (i) a stilbenoid isolated from extracts of *Lespedeza bicolor* mitigated PQ neurotoxicity in cell culture^59^; and (ii) that resveratrol alleviated oxidative stress in the brains of PQ-treated mice, resulting in improved motor function^60^. Based on these data and previous findings suggesting that resveratrol can penetrate the blood-brain barrier (BBB)^32, 61^, we infer that stilbenes or their metabolites may slow neurodegeneration in the brains of animals or humans exposed to PQ.

### A subset of individual ANC protect against PQ neurotoxicity

We found that four individual ANC – C3G, D3G, M3G, and C3So – protected against PQ neurotoxicity in our primary midbrain culture model. Three of these molecules (C3G, D3G, and M3G) were constituents of the BB and wild BB extracts, and C3G and D3G were also present in the BC extract. The lowest concentration of ANC at which we observed neuroprotective activity (0.05 µg/mL, in the case of D3G) was ∼900-fold higher than the concentration of <u>total</u> ANC in the diluted BB extract (at 0.01 µg/mL)^24^. These results substantiate the idea that synergistic interactions among multiple ANC in the BB extract produced an overall neuroprotective capacity that was substantially greater than the sum of the individual ANC activities^52^.

Our observation that the PB and hibiscus extracts failed to interfere with PQ neurotoxicity suggests that the malonyl and p-coumaryl glycoside derivatives of cyanidin that are abundant in the PB extract^39^, as well as the sambubioside derivatives of cyanidin and delphinidin in the hibiscus extract^38^, lack neuroprotective activity. Collectively, these data suggest that the acylated sugar moiety attached to the anthocyanidin core plays an important role in the ability of ANC to alleviate PQ neurotoxicity. The data presented here are likely relevant to neuroprotection *in vivo* given that intact anthocyanidin glycosides have been detected in the brains of rats and pigs fed with ANC- rich diets^62–64^.

### Different extracts have different neuroprotective activities against PQ versus rotenone

Previously, we reported that extracts prepared from BB, BC, GS, mulberry, and hibiscus (enriched in ANC, PAC, PA, and/or stilbenes – **Table 1**) protected against dopaminergic cell death caused by the PD-related neurotoxin, rotenone^24^. In the current study, we showed that all but one of these extracts (i.e., prepared from BB, BC, GS, or mulberry, but not hibiscus) also alleviated PQ neurotoxicity (**Table 2**)^24^. Moreover, we recently showed that an extract prepared from *Amelanchier arborea* interfered with dopaminergic cell death elicited by either PD-related insult^25^. Extracts that alleviate neurodegeneration caused by either insult are strong candidates for PD therapy because they could potentially antagonize different neurotoxic phenomena in the brains of individuals exposed to different PD-related toxins. In contrast, our observation that the PB extract failed to mitigate neurotoxicity elicited by either PQ or rotenone suggests that the abundant malonyl and p-coumaryl glycoside derivatives of cyanidin in this extract^39^ lack neuroprotective activity against either insult. Moreover, the fact that the hibiscus extract interfered with dopaminergic cell death in midbrain cultures exposed to rotenone but not PQ suggests that the sambubioside derivatives of cyanidin and delphinidin in this extract^38^ selectively activate responses that are protective against rotenone-mediated but not PQ-mediated neurotoxicity (discussed in more detail below).

**Table 2.**
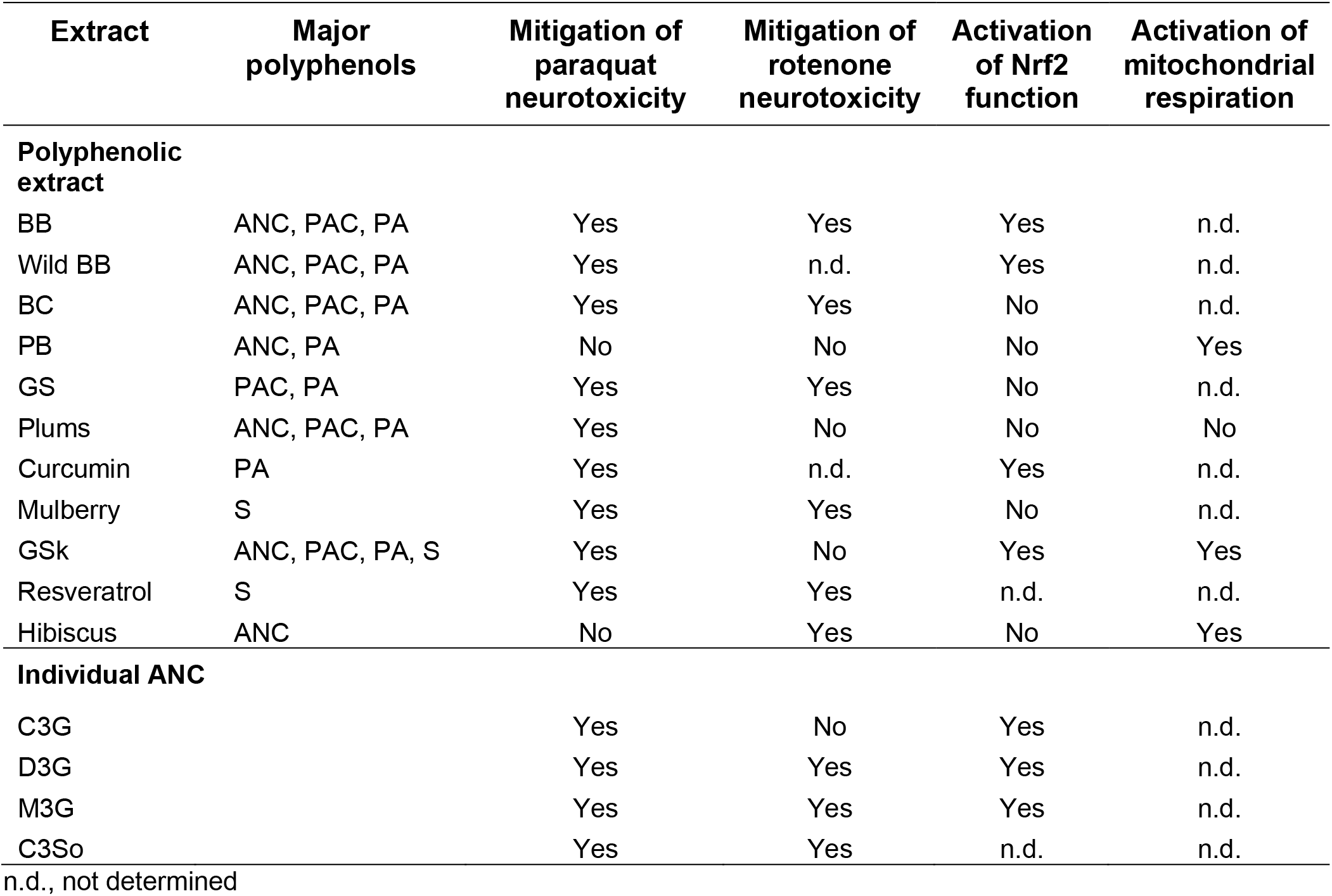
Summary of neuroprotective activities of botanical extracts and individual polyphenols.

| Extract | Major polyphenols | Mitigation of paraquat neurotoxicity | Mitigation of rotenone neurotoxicity | Activation of Nrf2 function | Activation of mitochondrial respiration |
| --- | --- | --- | --- | --- | --- |
| **Polyphenolic extract** |  |  |  |  |  |
| BB | ANC, PAC, PA | Yes | Yes | Yes | n.d. |
| Wild BB | ANC, PAC, PA | Yes | n.d. | Yes | n.d. |
| BC | ANC, PAC, PA | Yes | Yes | No | n.d. |
| PB | ANC, PA | No | No | No | Yes |
| GS | PAC, PA | Yes | Yes | No | n.d. |
| Plums | ANC, PAC, PA | Yes | No | No | No |
| Curcumin | PA | Yes | n.d. | Yes | n.d. |
| Mulberry | S | Yes | Yes | No | n.d. |
| GSk | ANC, PAC, PA, S | Yes | No | Yes | Yes |
| Resveratrol | S | Yes | Yes | n.d. | n.d. |
| Hibiscus | ANC | No | Yes | No | Yes |
| **Individual ANC** |  |  |  |  |  |
| C3G |  | Yes | No | Yes | n.d. |
| D3G |  | Yes | Yes | Yes | n.d. |
| M3G |  | Yes | Yes | Yes | n.d. |
| C3So |  | Yes | Yes | n.d. | n.d. |

n.d., not determined

### Different extracts have different abilities to activate Nrf2 signaling or rescue mitochondrial dysfunction

Our study revealed a correlation between the ability of an extract or individual ANC to interfere with PQ neurotoxicity and the activation of Nrf2-mediated transcription in astrocytes. All extracts and individual ANC that activated Nrf2-mediated transcription (BB, wild BB, and GSk extract; C3G, D3G, M3G, and curcumin) protected against PQ neurotoxicity (**Table 2**). Moreover, two extracts that failed to activate Nrf2-mediated transcription (prepared from PB or hibiscus) lacked neuroprotective activity against PQ-mediated dopaminergic cell death. From these data, we infer that activation of the Nrf2 pathway plays an important role in protection against PQ neurotoxicity. This result is consistent with our finding that Nrf2 expression is *sufficient* to alleviate PQ-mediated dopaminergic cell death in primary midbrain cultures (**Figure 6**). Moreover, other groups showed that neuronal cells are protected against PQ toxicity by treatment with a Nrf2 activator such as resveratrol^65^, sulforaphane^66^, or tert-butylhydroquinone^67^. Although BC, GS, plums, and mulberry extract failed to activate Nrf2-mediated transcription, they nevertheless alleviated PQ-induced neurotoxicity. This observation suggests that Nrf2 activation is not *necessary* for the protective effects of some extracts against PQ neurotoxicity.

Although C3G and D3G increased Nrf2 activity, C3Sa and D3Sa (the two ANC in the hibiscus extract) failed to do so. ANC are known to undergo a cycling mechanism that results in the formation of electrophilic quinones. ANC quinones can form adducts with the cysteine residues of Keap1, thereby activating the Nrf2 pathway^68, 69^. The inability of the hibiscus extract to activate Nrf2 signaling implies (as one possibility) that the sambubioside moiety of C3Sa or D3Sa is structurally incompatible with this mechanism, perhaps because of steric hindrance resulting from the larger size of sambubioside (a disaccharide) compared to the glucoside moiety of C3G or D3G.

We found that the hibiscus extract rescued rotenone-induced defects in mitochondrial respiration in galactose-conditioned N27 cells, an effect that could explain its protective activity against rotenone neurotoxicity in primary midbrain cultures^24^. In contrast, the plum extract failed to rescue rotenone-induced deficits in mitochondrial respiration, consistent with its inability to protect against rotenone neurotoxicity in midbrain cultures^24^. Because the PB and GSk extracts failed to protect against rotenone-induced dopaminergic neuron death in midbrain cultures^24^ despite rescuing rotenone-mediated defects in mitochondrial respiration in N27 cells, we infer that the ability of an extract to alleviate mitochondrial respiratory deficits is not sufficient for protection against rotenone neurotoxicity.

## Conclusions

In summary, our data show that extracts rich in ANC, PAC, and stilbenes alleviate neurotoxicity elicited by the PD-related insult, PQ. In addition, individual ANC were found to have protective activity against PQ neurotoxicity. These findings are consistent with epidemiological studies suggesting that a high intake of extracts rich in ANC and PAC is associated with a lower PD risk^36, 37^. A number of polyphenol-rich extracts or individual ANC exhibited neuroprotective activity against toxicity elicited by either PQ or rotenone. Nrf2 signaling was found to be sufficient for protection against PQ toxicity, whereas the ability to activate the Nrf2 pathway or ameliorate mitochondrial deficits is neither necessary nor sufficient for protection against toxicity elicited by rotenone. These insights into cellular mechanisms by which polyphenol-rich extracts protect against PQ- or rotenone-induced neurodegeneration advance our understanding of differences in the neurotoxic mechanisms of both PD-related insults. Importantly, extracts that alleviate neurotoxicity elicited by multiple PD stresses are attractive candidates for neuroprotection in humans because they may slow neurodegeneration in individuals with elevated PD risk resulting from exposure to a range of insults. In particular, the ANC-rich extracts and individual ANC shown here to be neuroprotective may interfere with dopaminergic neuron death *in vivo* based on evidence that ANC can penetrate the BBB as intact glycosides^62–64^.

## Acknowledgements

This work was supported by NIH grants R21 AG039718 and R03 DA027111 (J.-C. R), a Pilot Grant from the Purdue-UAB Botanicals Research Center (NIH P50 AT000477-06), and grants from the Branfman Family Foundation and Showalter Trust (J.-C.R.). The research described herein was conducted in a facility constructed with support from Research Facilities Improvement Program Grants number C06-14499 and C06-15480 from the National Center for Research Resources of the NIH. The authors would like to thank the members of the laboratory for important discussions and feedback.

---

**Supplementary Table 1.**
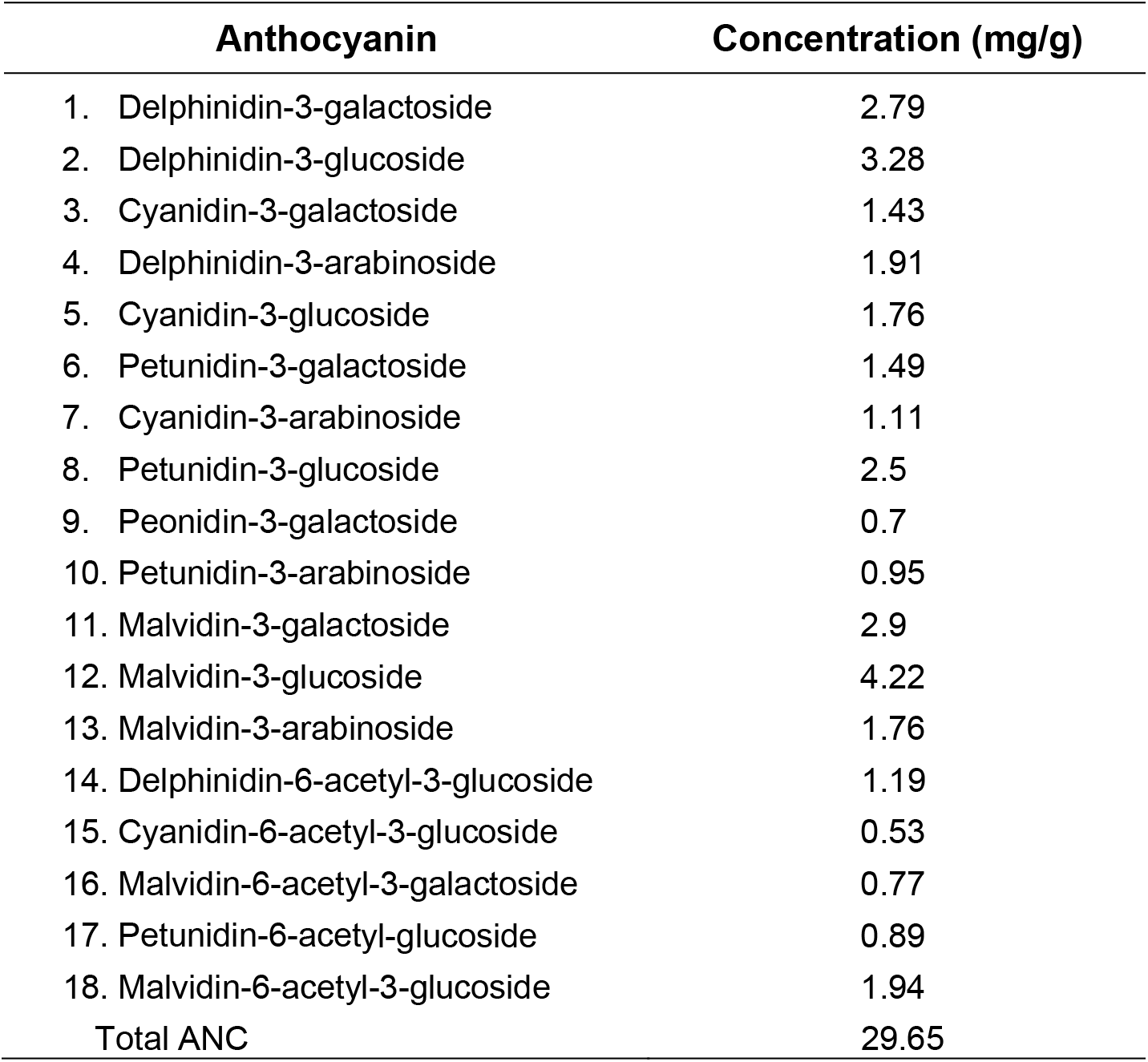
Anthocyanin composition of wild BB extract.

**Supplementary Table 2.**
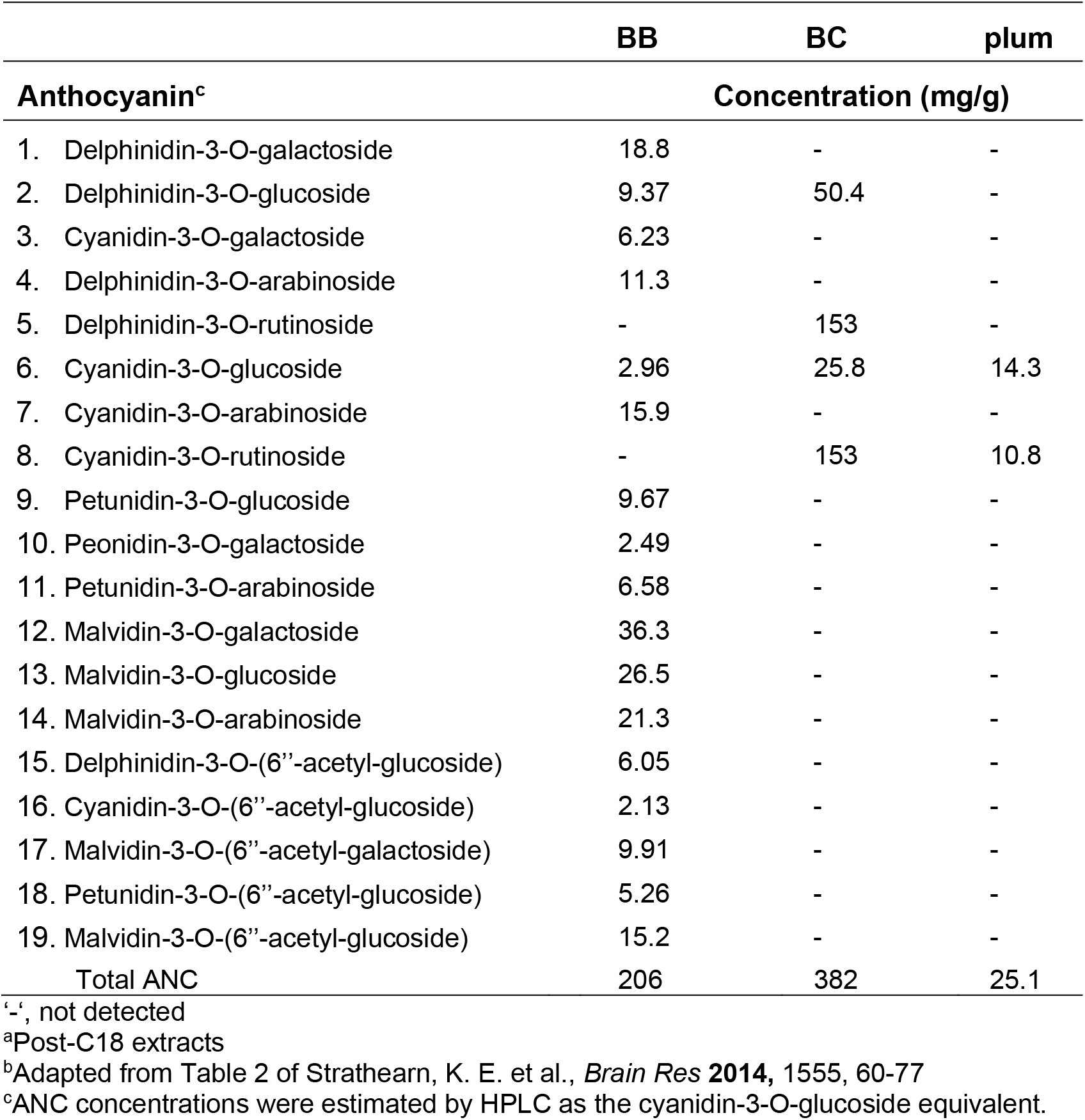
Anthocyanin composition of BB, BC, and plum extracts^a,b^.

**Supplementary Figure 1.**
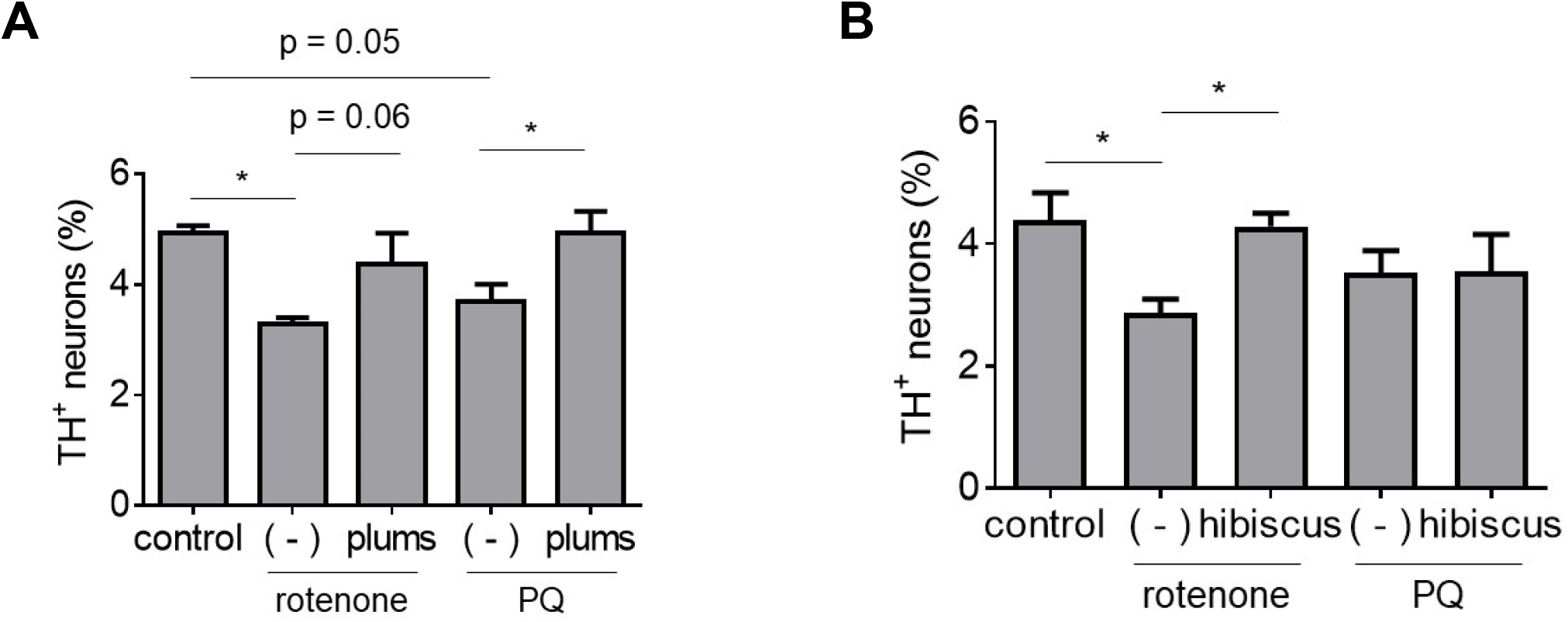
Comparison of protective activities of polyphenol-rich extracts against neurotoxicity elicited by rotenone or PQ. Primary midbrain cultures incubated in the absence or presence of extract prepared from plums (A) or hibiscus (B) for 72 h were exposed to rotenone (25 nM) or PQ (2.5 μM) in the absence or presence of extract for 24 h. Control cells were incubated in the absence of rotenone, PQ, or extract. The cells were stained with antibodies specific for MAP2 and TH and scored for relative dopaminergic cell viability. The data are presented as the mean ± SEM; *n* = 2 (B) or *n* = 3 (A), \**p*<0.05, square root transformation, one-way ANOVA with Tukey’s multiple comparisons post hoc test.

**Supplementary Figure 2.**
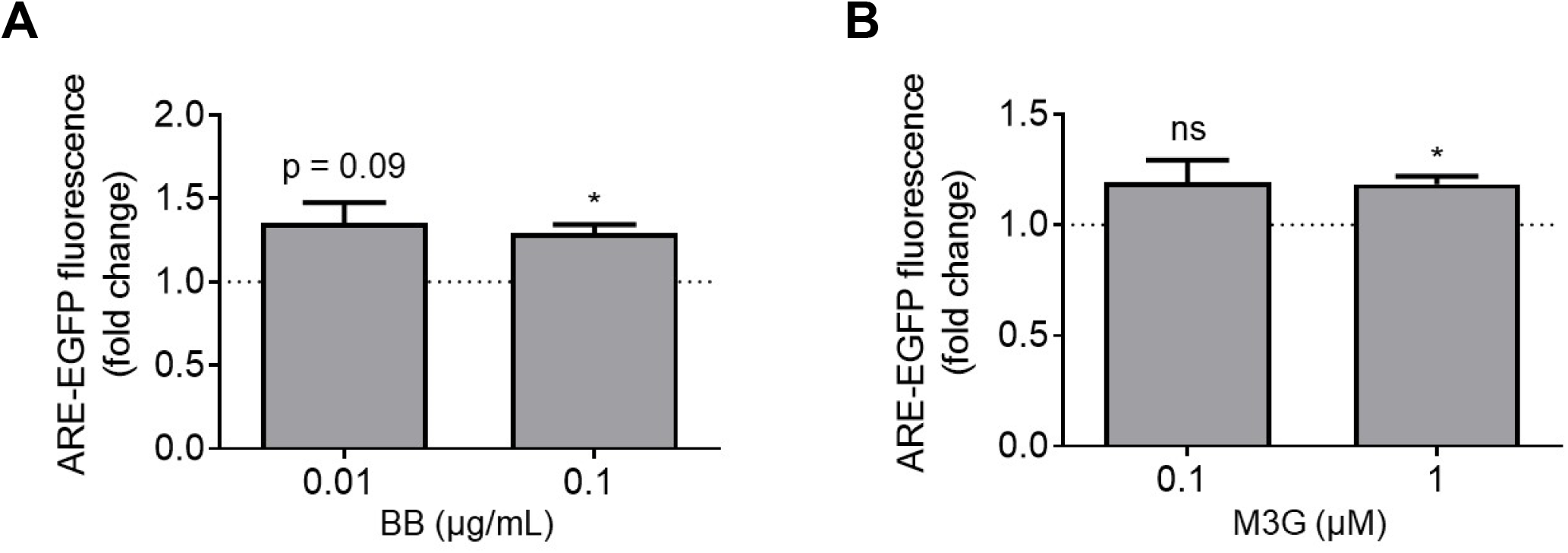
Evidence of an increase in Nrf2 transcriptional activity in human iPSC-derived astrocytes treated with BB extract or M3G. iCell astrocytes transduced with an ARE-EGFP reporter adenovirus for 48 h were incubated in the absence or presence of BB extract (A) or M3G (B) for 24 h. Control astrocytes were transduced with the reporter virus and incubated in the absence of extract or compound. The cells were imaged to determine the intracellular EGFP fluorescence intensity. The data are presented as the mean ± SEM; *n* = 3; \**p*<0.05 versus a predicted ratio of 1, log transformation followed by one-sample t-test (ns, not significant).

**Supplementary Figure 3.**
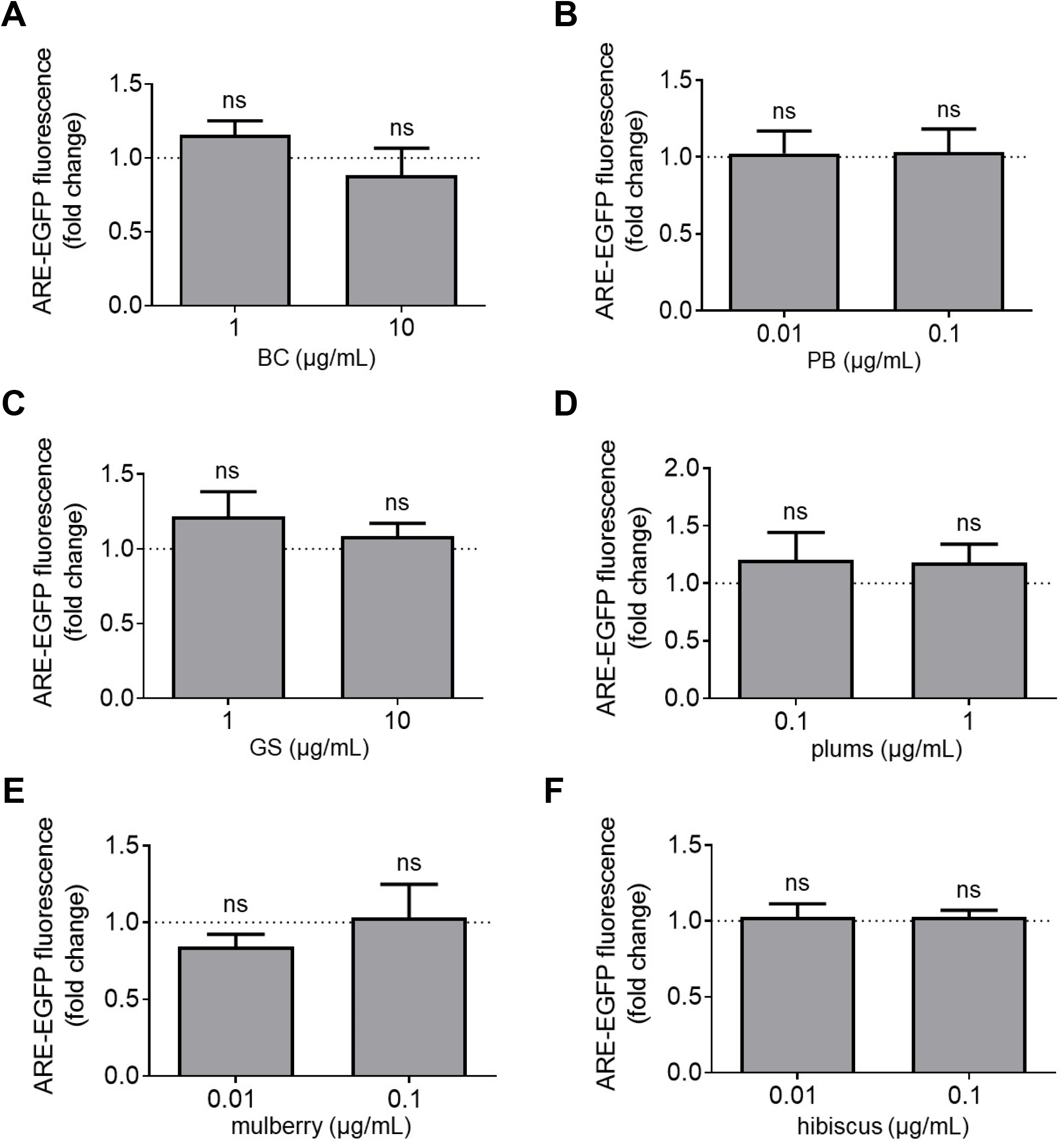
A subset of botanical extracts fail to activate Nrf2. Primary cortical astrocytes transduced with an ARE-EGFP reporter adenovirus for 48 h were incubated in the absence or presence of extract prepared from BC (A), PB (B), GS (C), plums (D), mulberry (E), or hibiscus (F) for 24 h. Control astrocytes were transduced with the reporter virus and incubated in the absence of extract. The cells were imaged to determine the intracellular EGFP fluorescence intensity. The data are presented as the mean ± SEM; *n* = 3 (A, C and D) or *n* = 4 (B) or *n* = 6 (E and F) (ns, not significant).

## REFERENCES

1. Gonzalez-Rodriguez, P.; Zampese, E.; Surmeier, D. J., Selective neuronal vulnerability in Parkinson’s disease. Prog Brain Res 2020, 252, 61–89.

2. Spillantini, M. G.; Schmidt, M. L.; Lee, V. M.-Y.; Trojanowski, J. Q.; Jakes, R.; Goedert, M., a-Synuclein in Lewy bodies. Nature 1997, 388, 839–840.

3. Estaun-Panzano, J.; Arotcarena, M. L.; Bezard, E., Monitoring α-synuclein aggregation. Neurobiol Dis 2023, 176, 105966.

4. Rochet, J. C.; Hay, B. A.; Guo, M., Molecular insights into Parkinson’s disease. Prog Mol Biol Transl Sci 2012, 107, 125–88.

5. Hernandez, D. G.; Reed, X.; Singleton, A. B., Genetics in Parkinson disease: Mendelian versus non-Mendelian inheritance. J Neurochem 2016, 139 Suppl 1, (Suppl 1), 59-74.

6. Huang, M.; Bargues-Carot, A.; Riaz, Z.; Wickham, H.; Zenitsky, G.; Jin, H.; Anantharam, V.; Kanthasamy, A.; Kanthasamy, A. G., Impact of Environmental Risk Factors on Mitochondrial Dysfunction, Neuroinflammation, Protein Misfolding, and Oxidative Stress in the Etiopathogenesis of Parkinson’s Disease. Int J Mol Sci 2022, 23, (18).

7. Conway, K. A.; Rochet, J.-C.; Bieganski, R. M.; Lansbury, P. T., Jr., Kinetic stabilization of the a-synuclein protofibril by a dopamine-a-synuclein adduct. Science 2001, 294, 1346–1349.

8. Ganguly, G.; Chakrabarti, S.; Chatterjee, U.; Saso, L., Proteinopathy, oxidative stress and mitochondrial dysfunction: cross talk in Alzheimer’s disease and Parkinson’s disease. Drug Des Devel Ther 2017, 11, 797–810.

9. Stykel, M. G.; Ryan, S. D., Nitrosative stress in Parkinson’s disease. NPJ Parkinsons Dis 2022, 8, (1), 104.

10. Tanner, C. M.; Kamel, F.; Ross, G. W.; Hoppin, J. A.; Goldman, S. M.; Korell, M.; Marras, C.; Bhudhikanok, G. S.; Kasten, M.; Chade, A. R.; Comyns, K.; Richards, M. B.; Meng, C.; Priestley, B.; Fernandez, H. H.; Cambi, F.; Umbach, D. M.; Blair, A.; Sandler, D. P.; Langston, J. W., Rotenone, Paraquat and Parkinson’s Disease. Environ Health Perspect 2011, 119, (6), 866-872.

11. Sherer, T. B.; Betarbet, R.; Testa, C. M.; Seo, B. B.; Richardson, J. R.; Kim, J. H.; Miller, G. W.; Yagi, T.; Matsuno-Yagi, A.; Greenamyre, J. T., Mechanism of toxicity in rotenone models of Parkinson’s disease. J. Neurosci. 2003, 23, (34), 10756–10764.

12. Betarbet, R.; Sherer, T. B.; MacKenzie, G.; Garcia-Osuna, M.; Panov, A. V.; Greenamyre, J. T., Chronic systemic pesticide exposure reproduces features of Parkinson’s disease. Nat. Neurosci. 2000, 3, 1301–1306.

13. Cannon, J. R.; Tapias, V.; Na, H. M.; Honick, A. S.; Drolet, R. E.; Greenamyre, J. T., A highly reproducible rotenone model of Parkinson’s disease. Neurobiol Dis 2009, 34, (2), 279–90.

14. Richardson, J. R.; Quan, Y.; Sherer, T. B.; Greenamyre, J. T.; Miller, G. W., Paraquat neurotoxicity is distinct from that of MPTP and rotenone. Toxicol Sci 2005, 88, (1), 193–201.

15. Bonneh-Barkay, D.; Langston, W. J.; Di Monte, D. A., Toxicity of redox cycling pesticides in primary mesencephalic cultures. Antioxid Redox Signal 2005, 7, (5-6), 649–53.

16. Cheatham, C. L.; Canipe, L. G., 3rd; Millsap, G.; Stegall, J. M.; Chai, S. C.; Sheppard, K. W.; Lila, M. A., Six-month intervention with wild blueberries improved speed of processing in mild cognitive decline: a double-blind, placebo-controlled, randomized clinical trial. Nutr Neurosci 2022, 1–15.

17. Cheatham, C. L.; Nieman, D. C.; Neilson, A. P.; Lila, M. A., Enhancing the Cognitive Effects of Flavonoids With Physical Activity: Is There a Case for the Gut Microbiome? Front Neurosci 2022, 16, 833202.

18. Lau, F. C.; Shukitt-Hale, B.; Joseph, J. A., Nutritional intervention in brain aging: reducing the effects of inflammation and oxidative stress. Subcell Biochem 2007, 42, 299–318.

19. Albarracin, S. L.; Stab, B.; Casas, Z.; Sutachan, J. J.; Samudio, I.; Gonzalez, J.; Gonzalo, L.; Capani, F.; Morales, L.; Barreto, G. E., Effects of natural antioxidants in neurodegenerative disease. Nutr Neurosci 2012, 15, (1), 1–9.

20. Akter, R.; Rahman, H.; Behl, T.; Chowdhury, M. A. R.; Manirujjaman, M.; Bulbul, I. J.; Elshenaw, S. E.; Tit, D. M.; Bungau, S., Prospective Role of Polyphenolic Compounds in the Treatment of Neurodegenerative Diseases. CNS Neurol Disord Drug Targets 2021, 20, (5), 430–450.

21. Levites, Y.; Weinreb, O.; Maor, G.; Youdim, M. B.; Mandel, S., Green tea polyphenol (-)- epigallocatechin-3-gallate prevents N-methyl-4-phenyl-1,2,3,6-tetrahydropyridine-induced dopaminergic neurodegeneration. J Neurochem 2001, 78, (5), 1073-82.

22. Guo, S.; Yan, J.; Yang, T.; Yang, X.; Bezard, E.; Zhao, B., Protective effects of green tea polyphenols in the 6-OHDA rat model of Parkinson’s disease through inhibition of ROS-NO pathway. Biol Psychiatry 2007, 62, (12), 1353–62.

23. Wang, Y.; Wu, S.; Li, Q.; Lang, W.; Li, W.; Jiang, X.; Wan, Z.; Chen, J.; Wang, H., Epigallocatechin-3-gallate: A phytochemical as a promising drug candidate for the treatment of Parkinson’s disease. Front Pharmacol 2022, 13, 977521.

24. Strathearn, K. E.; Yousef, G. G.; Grace, M. H.; Roy, S. L.; Tambe, M. A.; Ferruzzi, M. G.; Wu, Q. L.; Simon, J. E.; Ann Lila, M.; Rochet, J. C., Neuroprotective effects of anthocyanin- and proanthocyanidin-rich extracts in cellular models of Parkinson’s disease. Brain Res 2014, 1555, 60–77.

25. de Rus Jacquet, A.; Tambe, M. A.; Ma, S. Y.; McCabe, G. P.; Vest, J. H. C.; Rochet, J. C., Pikuni-Blackfeet traditional medicine: Neuroprotective activities of medicinal plants used to treat Parkinson’s disease-related symptoms. J Ethnopharmacol 2017, 206, 393–407.

26. de Rus Jacquet, A.; Timmers, M.; Ma, S. Y.; Thieme, A.; McCabe, G. P.; Vest, J. H. C.; Lila, M. A.; Rochet, J. C., Lumbee traditional medicine: Neuroprotective activities of medicinal plants used to treat Parkinson’s disease-related symptoms. J Ethnopharmacol 2017, 206, 408–425.

27. Winter, A. N.; Bickford, P. C., Anthocyanins and Their Metabolites as Therapeutic Agents for Neurodegenerative Disease. Antioxidants (Basel) 2019, 8, (9).

28. Jung, U. J.; Kim, S. R., Beneficial Effects of Flavonoids Against Parkinson’s Disease. J Med Food 2018, 21, (5), 421–432.

29. de Rus Jacquet, A.; Ambaw, A.; Tambe, M. A.; Ma, S. Y.; Timmers, M.; Grace, M. H.; Wu, Q. L.; Simon, J. E.; McCabe, G. P.; Lila, M. A.; Shi, R.; Rochet, J. C., Neuroprotective mechanisms of red clover and soy isoflavones in Parkinson’s disease models. Food Funct 2021, 12, (23), 11987–12007.

30. Sheta, R.; Teixeira, M.; Idi, W.; Pierre, M.; de Rus Jacquet, A.; Emond, V.; Zorca, C. E.; Vanderperre, B.; Durcan, T. M.; Fon, E. A.; Calon, F.; Chahine, M.; Oueslati, A., Combining NGN2 programming and dopaminergic patterning for a rapid and efficient generation of hiPSC-derived midbrain neurons. Sci Rep 2022, 12, (1), 17176.

31. Nebrisi, E. E., Neuroprotective Activities of Curcumin in Parkinson’s Disease: A Review of the Literature. Int J Mol Sci 2021, 22, (20).

32. Dos Santos, M. G.; Schimith, L. E.; André-Miral, C.; Muccillo-Baisch, A. L.; Arbo, B. D.; Hort, M. A., Neuroprotective Effects of Resveratrol in In vivo and In vitro Experimental Models of Parkinson’s Disease: a Systematic Review. Neurotox Res 2022, 40, (1), 319–345.

33. Ramassamy, C., Emerging role of polyphenolic compounds in the treatment of neurodegenerative diseases: a review of their intracellular targets. Eur J Pharmacol 2006, 545, (1), 51–64.

34. Chen, B.; Zhao, J.; Zhang, R.; Zhang, L.; Zhang, Q.; Yang, H.; An, J., Neuroprotective effects of natural compounds on neurotoxin-induced oxidative stress and cell apoptosis. Nutr Neurosci 2022, 25, (5), 1078–1099.

35. Limanaqi, F.; Biagioni, F.; Mastroiacovo, F.; Polzella, M.; Lazzeri, G.; Fornai, F., Merging the Multi-Target Effects of Phytochemicals in Neurodegeneration: From Oxidative Stress to Protein Aggregation and Inflammation. Antioxidants (Basel) 2020, 9, (10).

36. Gao, X.; Cassidy, A.; Schwarzschild, M. A.; Rimm, E. B.; Ascherio, A., Habitual intake of dietary flavonoids and risk of Parkinson disease. Neurology 2012, 78, (15), 1138–45.

37. Talebi, S.; Ghoreishy, S. M.; Jayedi, A.; Travica, N.; Mohammadi, H., Dietary Antioxidants and Risk of Parkinson’s Disease: A Systematic Review and Dose-Response Meta-analysis of Observational Studies. Adv Nutr 2022, 13, (5), 1493–1504.

38. Juliani, H. R.; Welch, C. R.; Wu, Q.; Diouf, B.; Malainy, D.; Simon, J. E., Chemistry and quality of Hibiscus (Hibiscus sabdariffa) for developing the natural-product industry in Senegal. J Food Sci 2009, 74, (2), S113–21.

39. Phippen, W. B.; Simon, J. E., Anthocyanins in Basil (Ocimum basilicum L.). J. Agric. Food Chem. 1998, 46, 1734–1738.

40. Wu, Q.; Wang, M.; Simon, J. E., Determination of proanthocyanidins in fresh grapes and grape products using liquid chromatography with mass spectrometric detection. Rapid Commun Mass Spectrom 2005, 19, (14), 2062–8.

41. Alam, J.; Stewart, D.; Touchard, C.; Boinapally, S.; Choi, A. M.; Cook, J. L., Nrf2, a Cap’n’Collar transcription factor, regulates induction of the heme oxygenase-1 gene. J Biol Chem 1999, 274, (37), 26071–8.

42. Alam, J.; Wicks, C.; Stewart, D.; Gong, P.; Touchard, C.; Otterbein, S.; Choi, A. M.; Burow, M. E.; Tou, J., Mechanism of heme oxygenase-1 gene activation by cadmium in MCF-7 mammary epithelial cells. Role of p38 kinase and Nrf2 transcription factor. J Biol Chem 2000, 275, (36), 27694-702.

43. Li, N.; Venkatesan, M. I.; Miguel, A.; Kaplan, R.; Gujuluva, C.; Alam, J.; Nel, A., Induction of heme oxygenase-1 expression in macrophages by diesel exhaust particle chemicals and quinones via the antioxidant-responsive element. J Immunol 2000, 165, (6), 3393–401.

44. Heurtaux, T.; Bouvier, D. S.; Benani, A.; Helgueta Romero, S.; Frauenknecht, K. B. M.; Mittelbronn, M.; Sinkkonen, L., Normal and Pathological NRF2 Signalling in the Central Nervous System. Antioxidants (Basel) 2022, 11, (8).

45. Bellavite, P., Neuroprotective Potentials of Flavonoids: Experimental Studies and Mechanisms of Action. Antioxidants (Basel) 2023, 12, (2).

46. Moratilla-Rivera, I.; Sánchez, M.; Valdés-González, J. A.; Gómez-Serranillos, M. P., Natural Products as Modulators of Nrf2 Signaling Pathway in Neuroprotection. Int J Mol Sci 2023, 24, (4).

47. Vargas, M. R.; Johnson, J. A., The Nrf2-ARE cytoprotective pathway in astrocytes. Expert Rev Mol Med 2009, 11, e17.

48. Bell, K. F.; Al-Mubarak, B.; Martel, M. A.; McKay, S.; Wheelan, N.; Hasel, P.; Márkus, N. M.; Baxter, P.; Deighton, R. F.; Serio, A.; Bilican, B.; Chowdhry, S.; Meakin, P. J.; Ashford, M. L.; Wyllie, D. J.; Scannevin, R. H.; Chandran, S.; Hayes, J. D.; Hardingham, G. E., Neuronal development is promoted by weakened intrinsic antioxidant defences due to epigenetic repression of Nrf2. Nat Commun 2015, 6, 7066.

49. Millichap, L. E.; Damiani, E.; Tiano, L.; Hargreaves, I. P., Targetable Pathways for Alleviating Mitochondrial Dysfunction in Neurodegeneration of Metabolic and Non-Metabolic Diseases. Int J Mol Sci 2021, 22, (21).

50. Marroquin, L. D.; Hynes, J.; Dykens, J. A.; Jamieson, J. D.; Will, Y., Circumventing the Crabtree effect: replacing media glucose with galactose increases susceptibility of HepG2 cells to mitochondrial toxicants. Toxicol Sci 2007, 97, (2), 539–47.

51. Case, A. J.; Agraz, D.; Ahmad, I. M.; Zimmerman, M. C., Low-Dose Aronia melanocarpa Concentrate Attenuates Paraquat-Induced Neurotoxicity. Oxid Med Cell Longev 2016, 2016, 5296271.

52. Williamson, E. M., Synergy and other interactions in phytomedicines. Phytomedicine 2001, 8, (5), 401–9.

53. Ortega-Arellano, H. F.; Jimenez-Del-Rio, M.; Velez-Pardo, C., Neuroprotective Effects of Methanolic Extract of Avocado Persea americana (var. Colinred) Peel on Paraquat-Induced Locomotor Impairment, Lipid Peroxidation and Shortage of Life Span in Transgenic knockdown Parkin Drosophila melanogaster. Neurochem Res 2019, 44, (8), 1986–1998.

54. Hou, R. R.; Chen, J. Z.; Chen, H.; Kang, X. G.; Li, M. G.; Wang, B. R., Neuroprotective effects of (-)-epigallocatechin-3-gallate (EGCG) on paraquat-induced apoptosis in PC12 cells. Cell Biol Int 2008, 32, (1), 22–30.

55. Ortega-Arellano, H. F.; Jimenez-Del-Rio, M.; Velez-Pardo, C., Life span and locomotor activity modification by glucose and polyphenols in Drosophila melanogaster chronically exposed to oxidative stress-stimuli: implications in Parkinson’s disease. Neurochem Res 2011, 36, (6), 1073–86.

56. Choi, J. Y.; Park, C. S.; Kim, D. J.; Cho, M. H.; Jin, B. K.; Pie, J. E.; Chung, W. G., Prevention of nitric oxide-mediated 1-methyl-4-phenyl-1,2,3,6-tetrahydropyridine-induced Parkinson’s disease in mice by tea phenolic epigallocatechin 3-gallate. Neurotoxicology 2002, 23, (3), 367-74.

57. Abd El Mohsen, M. M.; Kuhnle, G.; Rechner, A. R.; Schroeter, H.; Rose, S.; Jenner, P.; Rice-Evans, C. A., Uptake and metabolism of epicatechin and its access to the brain after oral ingestion. Free Radic Biol Med 2002, 33, (12), 1693–702.

58. Wang, J.; Ferruzzi, M. G.; Ho, L.; Blount, J.; Janle, E. M.; Gong, B.; Pan, Y.; Gowda, G. A.; Raftery, D.; Arrieta-Cruz, I.; Sharma, V.; Cooper, B.; Lobo, J.; Simon, J. E.; Zhang, C.; Cheng, A.; Qian, X.; Ono, K.; Teplow, D. B.; Pavlides, C.; Dixon, R. A.; Pasinetti, G. M., Brain-targeted proanthocyanidin metabolites for Alzheimer’s disease treatment. J Neurosci 2012, 32, (15), 5144–50.

59. Tarbeeva, D. V.; Pislyagin, E. A.; Menchinskaya, E. S.; Berdyshev, D. V.; Kalinovskiy, A. I.; Grigorchuk, V. P.; Mishchenko, N. P.; Aminin, D. L.; Fedoreyev, S. A., Polyphenolic Compounds from Lespedeza bicolor Protect Neuronal Cells from Oxidative Stress. Antioxidants (Basel) 2022, 11, (4).

60. Satpute, R. M.; Pawar, P. P.; Puttewar, S.; Sawale, S. D.; Ambhore, P. D., Effect of resveratrol and tetracycline on the subacute paraquat toxicity in mice. Hum Exp Toxicol 2017, 36, (12), 1303–1314.

61. Wang, J.; Tang, C.; Ferruzzi, M. G.; Gong, B.; Song, B. J.; Janle, E. M.; Chen, T. Y.; Cooper, B.; Varghese, M.; Cheng, A.; Freire, D.; Bilski, A.; Roman, J.; Nguyen, T.; Ho, L.; Talcott, S. T.; Simon, J. E.; Wu, Q.; Pasinetti, G. M., Role of standardized grape polyphenol preparation as a novel treatment to improve synaptic plasticity through attenuation of features of metabolic syndrome in a mouse model. Mol Nutr Food Res 2013, 57, (12), 2091–102.

62. Talavera, S.; Felgines, C.; Texier, O.; Besson, C.; Gil-Izquierdo, A.; Lamaison, J. L.; Remesy, C., Anthocyanin metabolism in rats and their distribution to digestive area, kidney, and brain. J Agric Food Chem 2005, 53, (10), 3902–8.

63. Milbury, P. E.; Kalt, W., Xenobiotic metabolism and berry flavonoid transport across the blood-brain barrier. J Agric Food Chem 2010, 58, (7), 3950–6.

64. Ho, L.; Ferruzzi, M. G.; Janle, E. M.; Wang, J.; Gong, B.; Chen, T. Y.; Lobo, J.; Cooper, B.; Wu, Q. L.; Talcott, S. T.; Percival, S. S.; Simon, J. E.; Pasinetti, G. M., Identification of brain-targeted bioactive dietary quercetin-3-O-glucuronide as a novel intervention for Alzheimer’s disease. Faseb J 2013, 27, (2), 769–81.

65. Zhang, L.; Dong, M. N.; Deng, J.; Zhang, C. H.; Liu, M. W., Resveratrol exhibits neuroprotection against paraquat-induced PC12 cells via heme oxygenase 1 upregulation by decreasing MiR-136-5p expression. Bioengineered 2022, 13, (3), 7065–7081.

66. Niso-Santano, M.; González-Polo, R. A.; Bravo-San Pedro, J. M.; Gómez-Sánchez, R.; Lastres-Becker, I.; Ortiz-Ortiz, M. A.; Soler, G.; Morán, J. M.; Cuadrado, A.; Fuentes, J. M., Activation of apoptosis signal-regulating kinase 1 is a key factor in paraquat-induced cell death: modulation by the Nrf2/Trx axis. Free Radic Biol Med 2010, 48, (10), 1370–81.

67. Li, H.; Wu, S.; Wang, Z.; Lin, W.; Zhang, C.; Huang, B., Neuroprotective effects of tert-butylhydroquinone on paraquat-induced dopaminergic cell degeneration in C57BL/6 mice and in PC12 cells. Arch Toxicol 2012, 86, (11), 1729–40.

68. Erlank, H.; Elmann, A.; Kohen, R.; Kanner, J., Polyphenols activate Nrf2 in astrocytes via H2O2, semiquinones, and quinones. Free Radic Biol Med 2011, 51, (12), 2319–27.

69. Jacob, J. K.; Tiwari, K.; Correa-Betanzo, J.; Misran, A.; Chandrasekaran, R.; Paliyath, G., Biochemical basis for functional ingredient design from fruits. Annu Rev Food Sci Technol 2012, 3, 79–104.

