## Supplementary Tables and Figures for "Protective effects of polyphenol-rich extracts against neurotoxicity elicited by paraquat or rotenone in cellular models of Parkinson’s disease"

**Supplementary Table 1. Anthocyanin composition of wild BB extract**

| Anthocyanin | Concentration (mg/g) |
|---|---|
| 1. Delphinidin-3-galactoside | 2.79 |
| 2. Delphinidin-3-glucoside | 3.28 |
| 3. Cyanidin-3-galactoside | 1.43 |
| 4. Delphinidin-3-arabinoside | 1.91 |
| 5. Cyanidin-3-glucoside | 1.76 |
| 6. Petunidin-3-galactoside | 1.49 |
| 7. Cyanidin-3-arabinoside | 1.11 |
| 8. Petunidin-3-glucoside | 2.5 |
| 9. Peonidin-3-galactoside | 0.7 |
| 10. Petunidin-3-arabinoside | 0.95 |
| 11. Malvidin-3-galactoside | 2.9 |
| 12. Malvidin-3-glucoside | 4.22 |
| 13. Malvidin-3-arabinoside | 1.76 |
| 14. Delphinidin-6-acetyl-3-glucoside | 1.19 |
| 15. Cyanidin-6-acetyl-3-glucoside | 0.53 |
| 16. Malvidin-6-acetyl-3-galactoside | 0.77 |
| 17. Petunidin-6-acetyl-glucoside | 0.89 |
| 18. Malvidin-6-acetyl-3-glucoside | 1.94 |
| Total ANC | 29.65 |

**Supplementary Table 2. Anthocyanin composition of BB, BC, and plum extracts[a,b]**

| | BB | BC | plum |
|---|---|---|---|
| **Anthocyanin[c]** | | **Concentration (mg/g)** | |
| 1. Delphinidin-3-O-galactoside | 18.8 | - | - |
| 2. Delphinidin-3-O-glucoside | 9.37 | 50.4 | - |
| 3. Cyanidin-3-O-galactoside | 6.23 | - | - |
| 4. Delphinidin-3-O-arabinoside | 11.3 | - | - |
| 5. Delphinidin-3-O-rutinoside | - | 153 | - |
| 6. Cyanidin-3-O-glucoside | 2.96 | 25.8 | 14.3 |
| 7. Cyanidin-3-O-arabinoside | 15.9 | - | - |
| 8. Cyanidin-3-O-rutinoside | - | 153 | 10.8 |
| 9. Petunidin-3-O-glucoside | 9.67 | - | - |
| 10. Peonidin-3-O-galactoside | 2.49 | - | - |
| 11. Petunidin-3-O-arabinoside | 6.58 | - | - |
| 12. Malvidin-3-O-galactoside | 36.3 | - | - |
| 13. Malvidin-3-O-glucoside | 26.5 | - | - |
| 14. Malvidin-3-O-arabinoside | 21.3 | - | - |
| 15. Delphinidin-3-O-(6''-acetyl-glucoside) | 6.05 | - | - |
| 16. Cyanidin-3-O-(6''-acetyl-glucoside) | 2.13 | - | - |
| 17. Malvidin-3-O-(6''-acetyl-galactoside) | 9.91 | - | - |
| 18. Petunidin-3-O-(6''-acetyl-glucoside) | 5.26 | - | - |
| 19. Malvidin-3-O-(6''-acetyl-glucoside) | 15.2 | - | - |
| Total ANC | 206 | 382 | 25.1 |

'-', not detected
[a]Post-C18 extracts
[b]Adapted from Table 2 of Strathearn, K. E. et al., *Brain Res* **2014,** 1555, 60-77
[c]ANC concentrations were estimated by HPLC as the cyanidin-3-O-glucoside equivalent.

### Supplementary Figure 1

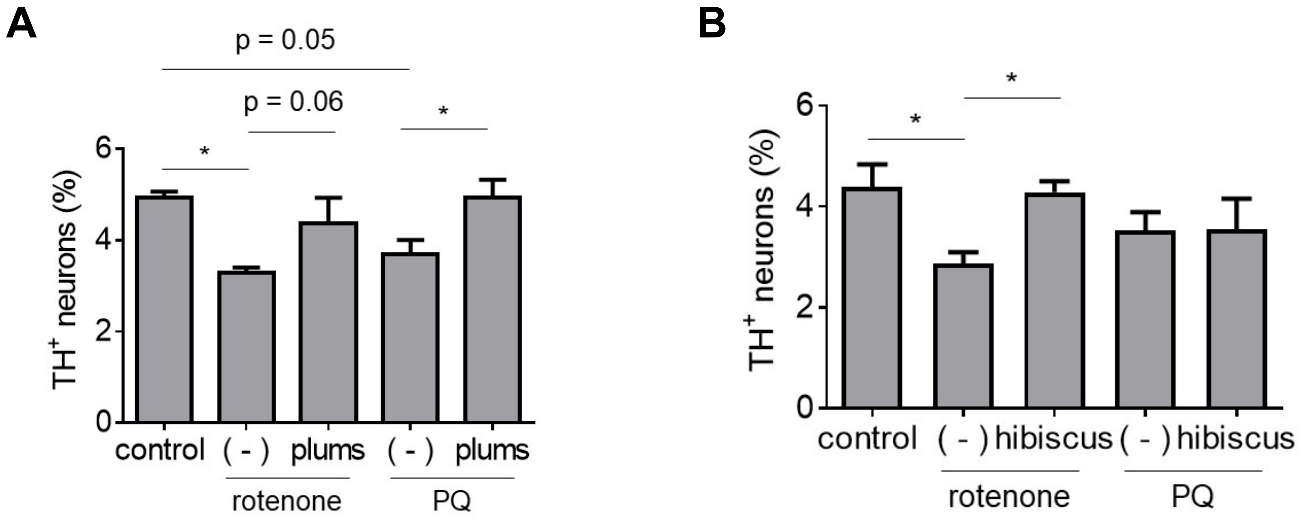


**Supplementary Figure 1. Comparison of protective activities of polyphenol-rich extracts against neurotoxicity elicited by rotenone or PQ.** Primary midbrain cultures incubated in the absence or presence of extract prepared from plums (A) or hibiscus (B) for 72 h were exposed to rotenone (25 nM) or PQ (2.5 μM) in the absence or presence of extract for 24 h. Control cells were incubated in the absence of rotenone, PQ, or extract. The cells were stained with antibodies specific for MAP2 and TH and scored for relative dopaminergic cell viability. The data are presented as the mean ± SEM; $n$ = 2 (B) or $n$ = 3 (A), *$p$<0.05, square root transformation, one-way ANOVA with Tukey's multiple comparisons post hoc test.

Supplementary Figure 2

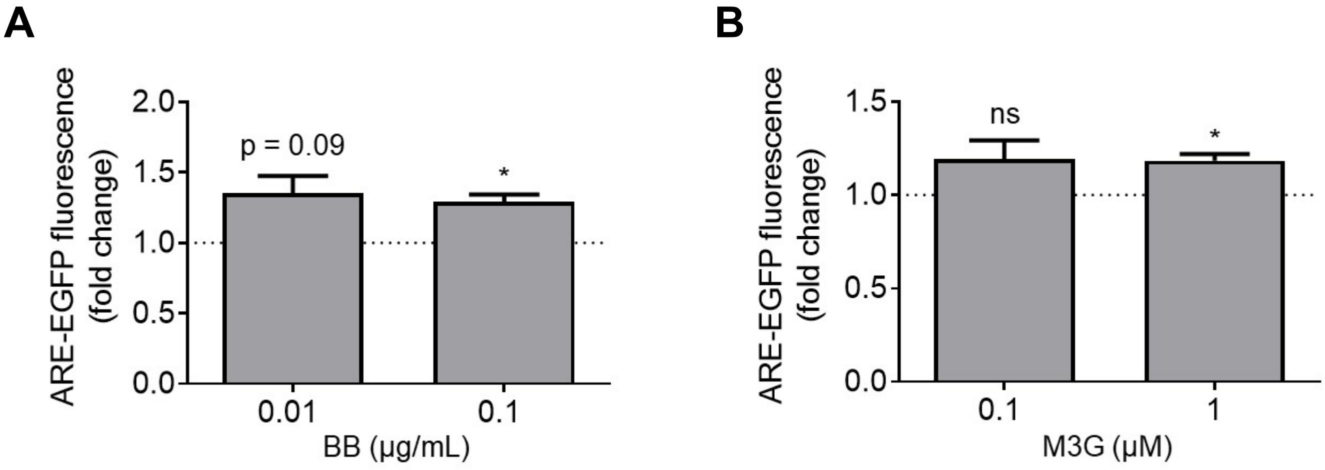


**Supplementary Figure 2. Evidence of an increase in Nrf2 transcriptional activity in human iPSC-derived astrocytes treated with BB extract or M3G.** iCell astrocytes transduced with an ARE-EGFP reporter adenovirus for 48 h were incubated in the absence or presence of BB extract (A) or M3G (B) for 24 h. Control astrocytes were transduced with the reporter virus and incubated in the absence of extract or compound. The cells were imaged to determine the intracellular EGFP fluorescence intensity. The data are presented as the mean ± SEM; $n = 3$; *$p<0.05$ versus a predicted ratio of 1, log transformation followed by one-sample t-test (ns, not significant).

### Supplementary Figure 3

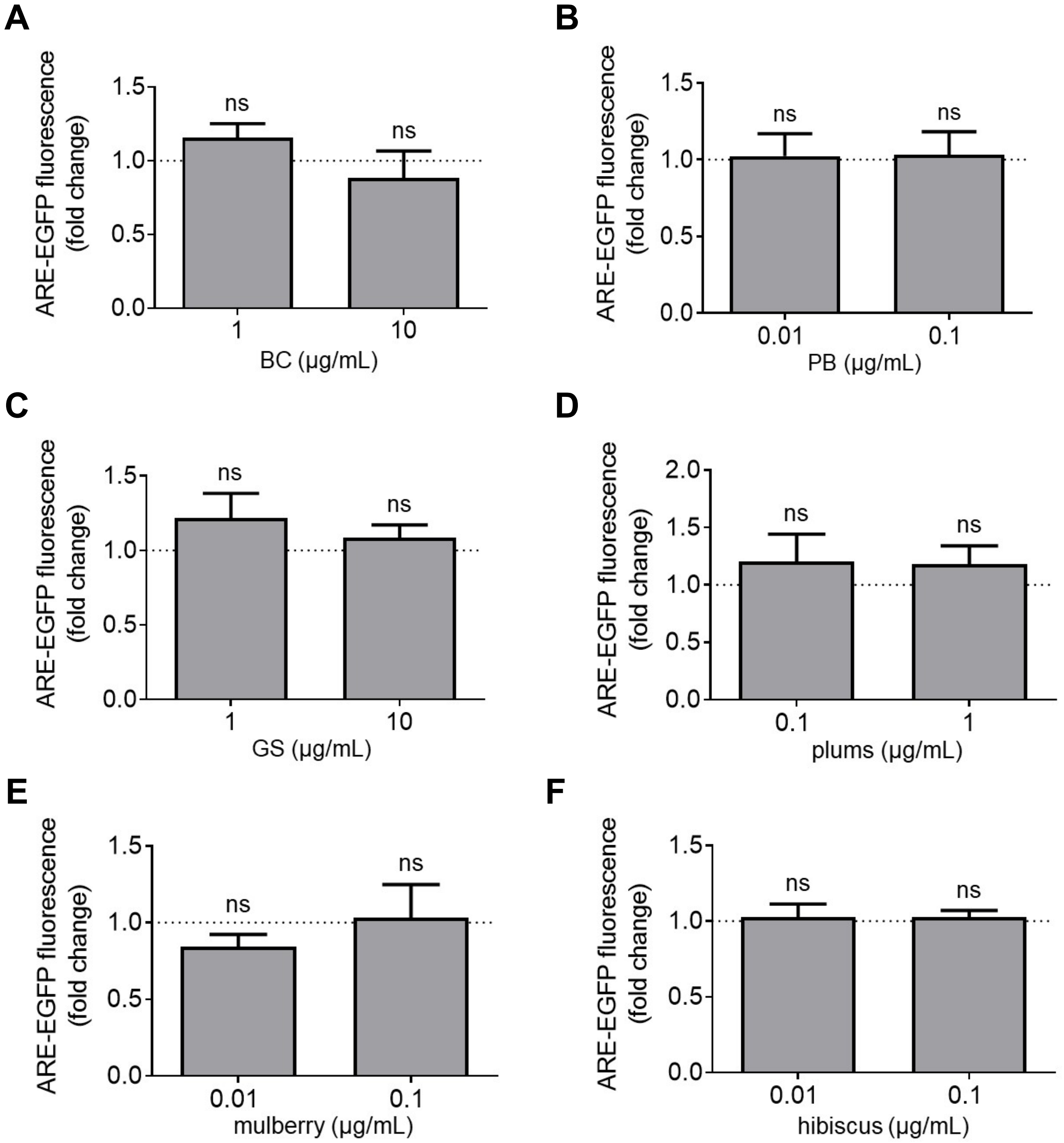


**Supplementary Figure 3. A subset of botanical extracts fail to activate Nrf2.** Primary cortical astrocytes transduced with an ARE-EGFP reporter adenovirus for 48 h were incubated in the absence or presence of extract prepared from BC (A), PB (B), GS (C), plums (D), mulberry (E), or hibiscus (F) for 24 h. Control astrocytes were transduced with the reporter virus and incubated in the absence of extract. The cells were imaged to determine the intracellular EGFP fluorescence intensity. The data are presented as the mean ± SEM; $n$ = 3 (A, C and D) or $n$ = 4 (B) or $n$ = 6 (E and F) (ns, not significant).
